# Haplotype-resolved reconstruction and functional interrogation of cancer karyotypes

**DOI:** 10.1101/2024.03.02.583108

**Authors:** Gregory J. Brunette, Richard W. Tourdot, Darawalee Wangsa, David Pellman, Cheng-Zhong Zhang

## Abstract

Genomic characterization has revealed widespread structural complexity in cancer karyotypes, however shotgun sequencing cannot resolve genomic rearrangements with chromosome-length continuity. Here, we describe a two-tiered approach to determine the segmental composition of rearranged chromosomes with haplotype resolution. First, we present *refLinker*, a new method for robust determination of chromosomal haplotypes using cancer Hi-C data. Second, we use haplotype-specific Hi-C contacts to determine the segmental structure of rearranged chromosomes. By contrast with existing methods for diploid haplotype inference, our approach is robust to the confounding effects of large-scale DNA deletions, duplications, and high-level amplification in cancer sequencing. Using this approach, we examine haplotype-specific expression changes on rearranged homologs and provide direct evidence for long-range transcriptional activation and repression associated with rearrangements of the inactive X chromosome (Xi). Together, these results reveal the significant transcriptional consequences of somatic Xi rearrangements, highlighting *refLinker*’s broad utility for studying the functional consequences of chromosomal rearrangements.

## Introduction

Cytogenetic analysis and DNA sequencing revealed widespread alterations to chromosome number and structure in cancer genomes, advancing our understanding of how specific chromosomal rearrangements contribute to disease [1], [2], [3]. Despite these advances, it remains challenging to relate nucleotide-level variation to the long-range structure of rearranged chromosomes. Shotgun sequencing can reveal DNA breakpoints at base-pair resolution, but has limitations for assembling these breakpoints into the larger context of rearranged chromosomes. On the other hand, cytogenetic karyotyping enables the direct visualization of large-scale chromosomal aberrations but is unable to precisely map genomic breakpoints. Neither shotgun sequencing nor cytogenetics can accurately determine the parental homolog origin of rearranged DNA segments, limiting our ability to investigate both how these alterations arise and their functional consequences (**Figure 1A**). Although these problems are partly addressed by long read sequencing [4], [5], [6], cancer genomes typically harbor large duplicated segments (1-10 Mb) with little genetic variation (< 1 variant per 100 kb), making it difficult or impossible for long reads (10-100 kb) to determine the breakpoints joining these segments.

**Figure 1.**
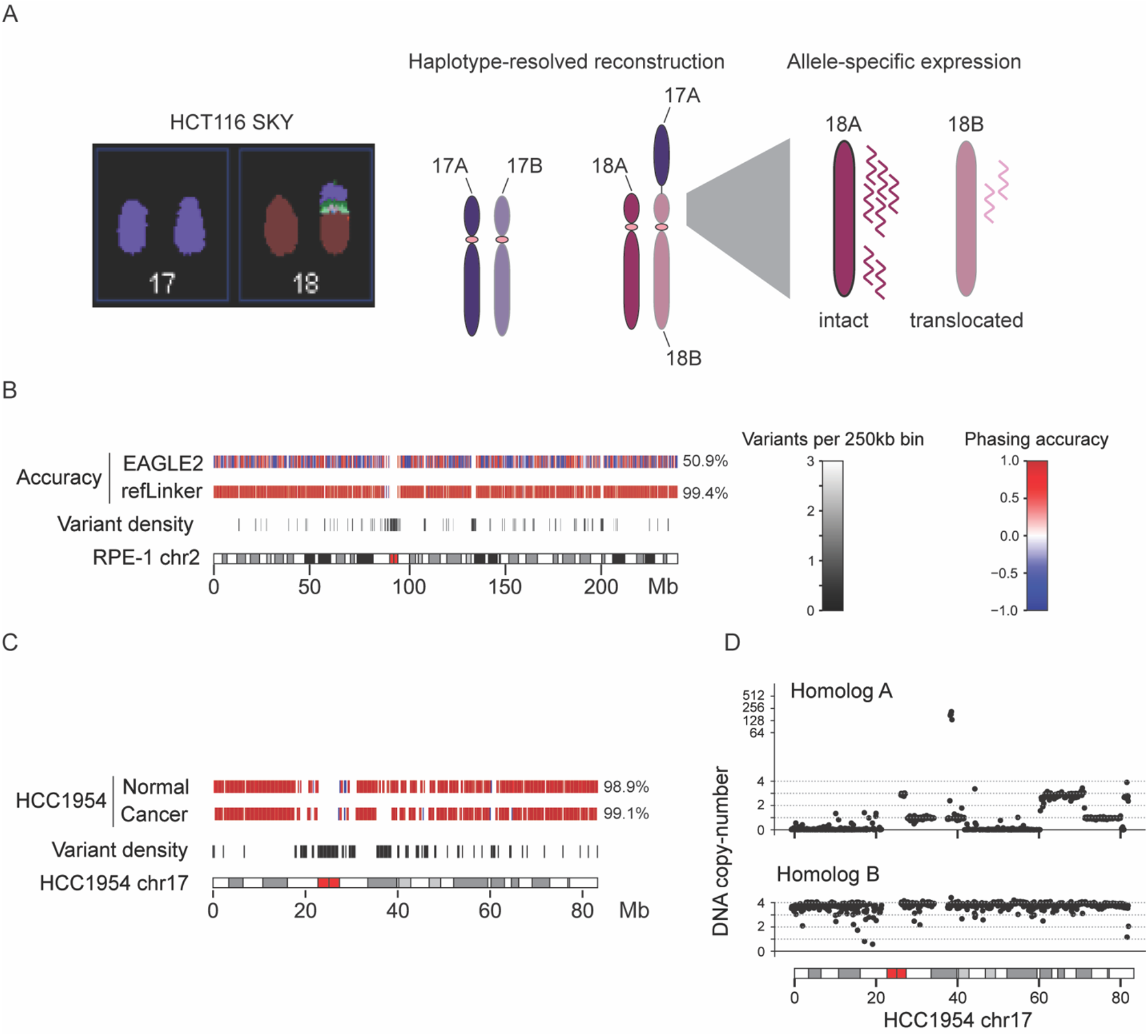
(**A**) Outline of haplotype-resolved karyotype analysis. Left: SKY imaging depicts intact and translocated chr17 and chr18 in HCT116. Center: Haplotype-resolved karyotype construction of HCT116 resolved long-range sequence identity of chromosome homologs and Right: enables direct functional comparison of intact versus translocated alleles. (**B**) Switching errors in statistical phasing are eliminated by *refLinker* to generate accurate whole-chromosome haplotypes. Top row, switching errors in the EAGLE2 haplotype are indicated by red-to-blue phasing switches (500 kb bins) on chr2 of the RPE-1 genome; the average phasing accuracy of 50.9% reflects roughly equal contributions of both parental haplotypes in the EAGLE2 solution. Second row, consistent phasing accuracy after corrections using Hi-C links by *refLinker*. Third row, variant density (250kb bins) showing gaps in the haplotype solution corresponding to variant-poor regions, including the centromere. (**C**) As in (**B**), except *refLinker* was run for HCC1954 matched normal (top track, 98.9% accuracy) and cancer data (second track, 99.1% accuracy). (**D**) DNA copy-number of fragmented (top) and intact (bottom) HCC1954 chr17 homologs.

The difficulties in resolving cancer genomic rearrangements highlight a fundamental problem in resolving the structure of non-haploid genomes: determining the genomic locations of homologous DNA sequences contained in either rearranged chromosomes or different parental homologs. This problem is partly addressed by Hi-C sequencing, which preserves long-range (>1 Mb) intrachromosomal (*cis*) contacts. Hi-C’s utility for chromosome-scale genome reconstruction has been demonstrated by several recent studies [7], [8], [9], [10], [11], [12]. However, in aneuploid cancer genomes, the density of *cis* Hi-C contacts is confounded by both chromosomal copy-number alterations and translocations. The prevalence of such alterations prevents reliable haplotype determination and genome construction using state-of-the-art methods that are designed for haploid or diploid genomes.

To overcome this technical challenge, we present *refLinker*, a method that can robustly resolve whole-chromosome haplotypes with >99% accuracy and encompassing >99.9% of common variants from Hi-C sequencing of aneuploid cancer cells. We demonstrate the robust performance of haplotype inference by *refLinker* in the presence of allelic imbalance and structural complexity, and its capability to resolve rearranged chromosomes harboring large segmental duplications, inversions, and focal amplifications.

As a specific application of *refLinker*’s capability for cancer genome construction, we determine the segmental structure of rearranged X chromosomes in breast cancer cell lines and provide direct evidence for long-range transcriptional changes associated with translocations between autosomes and the inactive X chromosome (Xi). Whereas previous studies suggested a limited role for the *cis*-regulatory X-inactivation center (XIC) outside embryonic development [13] [14], we identified both transcriptional re-activation of Xi genes following XIC removal and transcriptional suppression of autosomal genes that are translocated to the XIC. Together, these findings highlight the plasticity of X-chromosome inactivation after structural rearrangement and identify a multi-pronged source of transcriptional dysregulation in cancer development, highlighting *refLinker*’s broad utility for relating chromosome structural alterations to functional outcomes.

## Results

### refLinker enables robust chromosomal haplotype inference from Hi-C data

Determination of the haplotype phase (i.e., the genotypes at heterozygous sites in each parental chromosome) is essential for the assembly and functional interrogation of rearrangements occurring to each parental chromosome (**Figure 1A**). To determine the haplotype with chromosome-scale accuracy requires Hi-C sequencing; the capability of Hi-C sequencing for chromosomal haplotype inference has been demonstrated in several studies [15], [16], [17]. However, these methods are usually developed for haplotype inference of diploid genomes; the accuracy and robustness of these methods when applied to cancer genomes are often compromised by extensive allelic imbalance and chromosomal structural alterations, which disrupt the molecular linkage within parental homologs and prevent precise genome construction. It is also unclear whether chromosome-scale haplotype inference can be achieved with Hi-C data alone, or requires additional long-read or long-range data types.

We have developed *refLinker* (https://github.com/gbrunette/refLinker), a software tool that can generate whole-chromosome haplotype phase from Hi-C data in combination with statistical phasing. Statistical phasing can derive haplotype linkage between common polymorphic genotypes in an individual genome using linkage disequilibrium in the human population. The accuracy of statistical phasing is limited to 10^4^-10^5^ kb (**Figure S1A**,**B**). Starting from a statistical phasing haplotype, *refLinker* uses Hi-C contacts to iteratively correct long-range phasing errors, which appear as regions of discordant molecular linkage between opposite homologs (**Figure S1C, D**). This approach determines haplotype linkage with chromosome-length accuracy (**Figure 1B**). For a full description of *refLinker* and its implementation, see **SI: Approach**.

To evaluate *refLinker*’s performance, we first tested its accuracy in three diploid human genomes with ground truth haplotype information: NA12878, RPE-1, and HCC1954BL. See **Table S1** for Hi-C data sources. In all cases, *refLinker* resolved the haplotype phase of common variants at >99% accuracy (**Figure 1B**, RPE-1 chr2; **Table S2-5** all chromosomes). We further ran *refLinker* on downsampled Hi-C data of RPE-1 cells (245,700,771 reads, 29.5 million long-range contacts between loci separated by ≥1Mb). Our analysis demonstrated that *refLinker* consistently achieved >98% accuracy with ∼10 million long-range contacts (**Table S6**, 0.333× downsampling), and >90% accuracy with ∼5 million long-range contacts (**Table S7**, 0.167× downsampling). Together, these results demonstrated the sufficiency of Hi-C data alone for whole-chromosome haplotype inference in diploid genomes.

We then compared *refLinker*’s accuracy to *IntegratedPhasing* [17], the only comparable method for these data types. IntegratedPhasing and *refLinker* are built on different formalisms, leading to distinct approaches for both (1) the integration of haplotype linkage from statistical phasing and (2) the quantification of Hi-C linkage within homologs (**SI: Benchmarking IntegratedPhasing in diploid genomes**). By contrast to *refLinker*, we found that *IntegratedPhasing* does not ensure the long-range phasing accuracy that is required for haplotype-specific resolution of long-range rearrangements in cancer genomes. This was demonstrated by the presence of arm-level switching errors and lesser long-range accuracy of *IntegratedPhasing* relative to *refLinker* (**Figure S2**).

### refLinker enables direct determination of cancer haplotypes from Hi-C sequencing

To assess whether Hi-C data can reliably determine the haplotype phase of chromosomes in aneuploid genomes, we next performed haplotype inference in HCC1954, a highly aneuploid breast cancer genome, using *refLinker* and *IntegratedPhasing*.

HCC1954 is near-tetraploid and contains extensive copy-number alterations and structural rearrangements that disrupt parental homolog structure. Despite such complexity, we expect that the haplotype phase of each parental chromosome can still be reliably determined as long as recombination between different homologous chromosomes is rare [18] (such recombination will create spurious Hi-C linkage between different homologs). Having already determined the haplotype phase of parental chromosomes of HCC1954 cells in its matching lymphoblastoid cell line HCC1954BL (**Table S5**), we could assess whether the parental haplotype can also be determined from the cancer Hi-C reads alone.

We found that *refLinker* achieved similar accuracy using cancer Hi-C reads when compared to the haplotype determined directly from Hi-C of the matching germline genome (**Figure 1C**, shown for chr17, **Table S9**, all chromosomes), in spite of extensive allelic imbalance and copy-number alterations in the cancer genome (**Figure 1D**). (We note that haplotype inference in hemizygous regions requires knowledge of heterozygous genotypes in the parental genome.) Comparing *refLinker* to *IntegratedPhasing*, we observed that *IntegratedPhasing* generated frequent haplotype switching in regions of allelic copy-number imbalance (**Figure 2A**), loss-of-heterozygosity (**Figure 2B**), or high-level copy-number amplification (**Figure 2C**). These errors highlight the difference in the computational algorithms. Briefly, *IntegratedPhasing* and other methods separately consider four types of molecular linkage: maternal-maternal, paternal-paternal, or either combination of the two, assuming the maximum of these four classes reflects intra-chromosomal linkage (either maternal-maternal or paternal-paternal). In complex genomes, this scheme can lead to false haplotype assignments. For example, large segmental deletions decrease the number of *cis*

**Figure 2.**
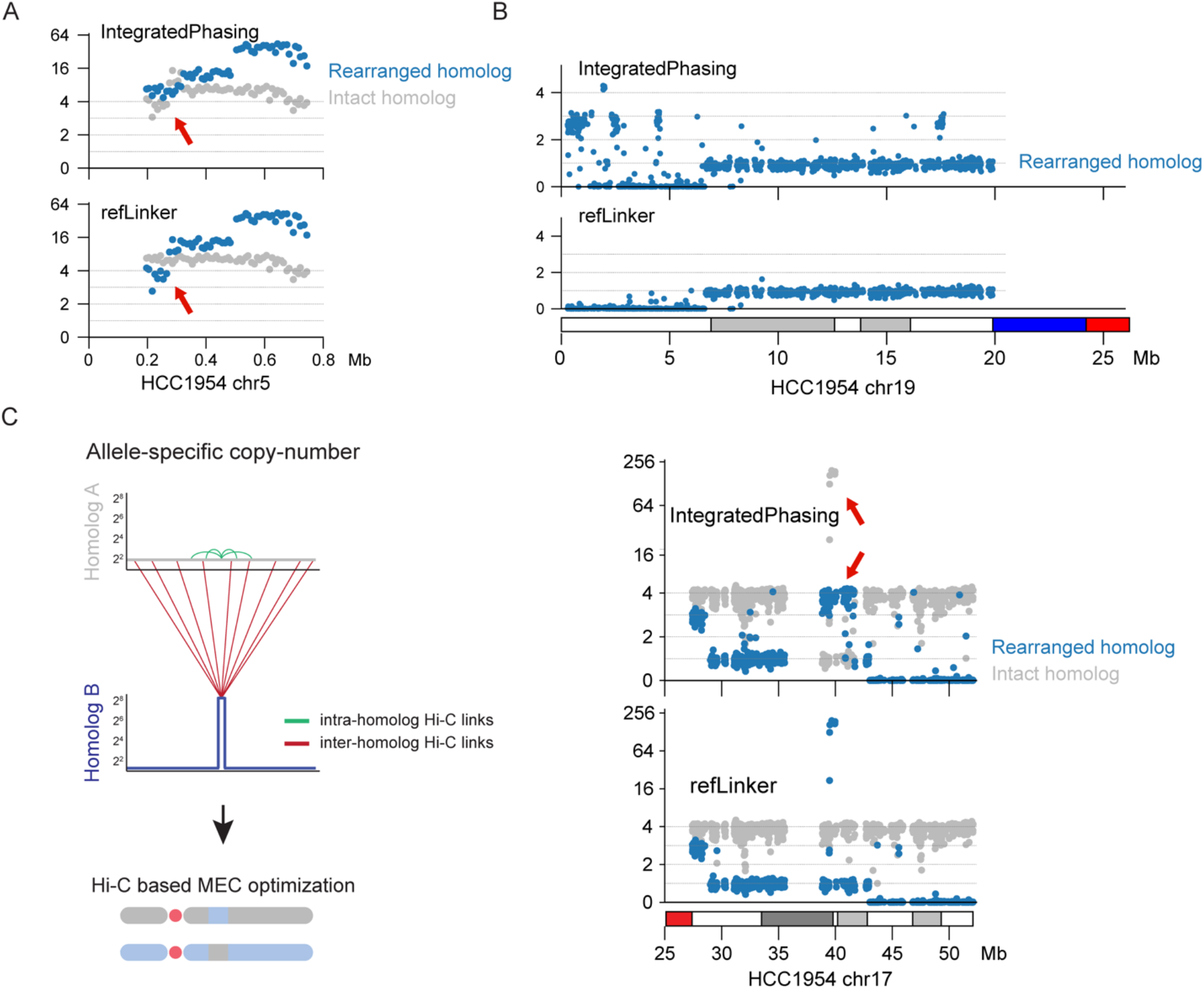
refLinker enables direct whole-chromosome haplotype determination from cancer Hi-C data. (**A-C**) Homolog-specific DNA copy-number was calculated for HCC1954 using IntegratedPhasing (top) and refLinker (bottom) haplotypes. Highlighted (red arrows) are examples of long-range phasing errors arising from (**A**) Allelic imbalance (**B**) Loss-of-heterozygosity (**C**) Focal amplification on different chromosomes of HCC1954. Left: schematic illustrating non-specific Hi-C interactions between chr17 amplicon and intact homolog.

Hi-C contacts within the retained segments. Likewise, high-level focal amplification can amplify *trans* Hi-C contacts between the amplified region and the opposite homolog. Together, these two alterations result in more *trans* Hi-C contacts between the amplified region and the opposite, intact homolog as shown in **Figure 2C** (left), leading to haplotype switching errors in *IntegratedPhasing*. By contrast, *refLinker* achieves the correct haplotype assignment in each of these cases because it jointly considers all four types of molecular linkage (**Figure S3**). These results highlight *refLinker*’s unique advantages for cancer genome reconstruction.

### Haplotype-specific reconstruction of simple and complex rearrangements

We next demonstrated the capability to determine the segmental structure of rearranged chromosomes using haplotype-specific Hi-C contacts. In contrast to previous studies using Hi-C for the determination of rearranged chromosome structure [19], [20], here the capability of chromosomal haplotype inference from *refLinker* enabled us to assemble the structure of rearranged chromosomes involving different parental homologs.

We analyzed the widely studied colorectal cancer cell line HCT116. HCT116 cells are near-diploid with three translocations visible by standard karyotyping: der(10)dup(10)(q?)t(10;16), der(16)t(8;16), and der(18)t(17;18) (**Figure 3A**) [21]. We determined the haplotype phase of parental chromosomes of HCT116 using publicly available whole-genome and Hi-C sequencing data [22], [23]. The haplotype phase contained 1,757,238 phased variants (**Supplementary Dataset 1**). The lower number of phased variants in comparison to diploid genomes is due to large regions of uniparental disomy on the p-arms of chr3, chr5, and chr7 (**Figure S4**). We note that the parental origin of the retained haplotype in these regions cannot be established as the same segments are present on both homologs for these three cases.

**Figure 3.**
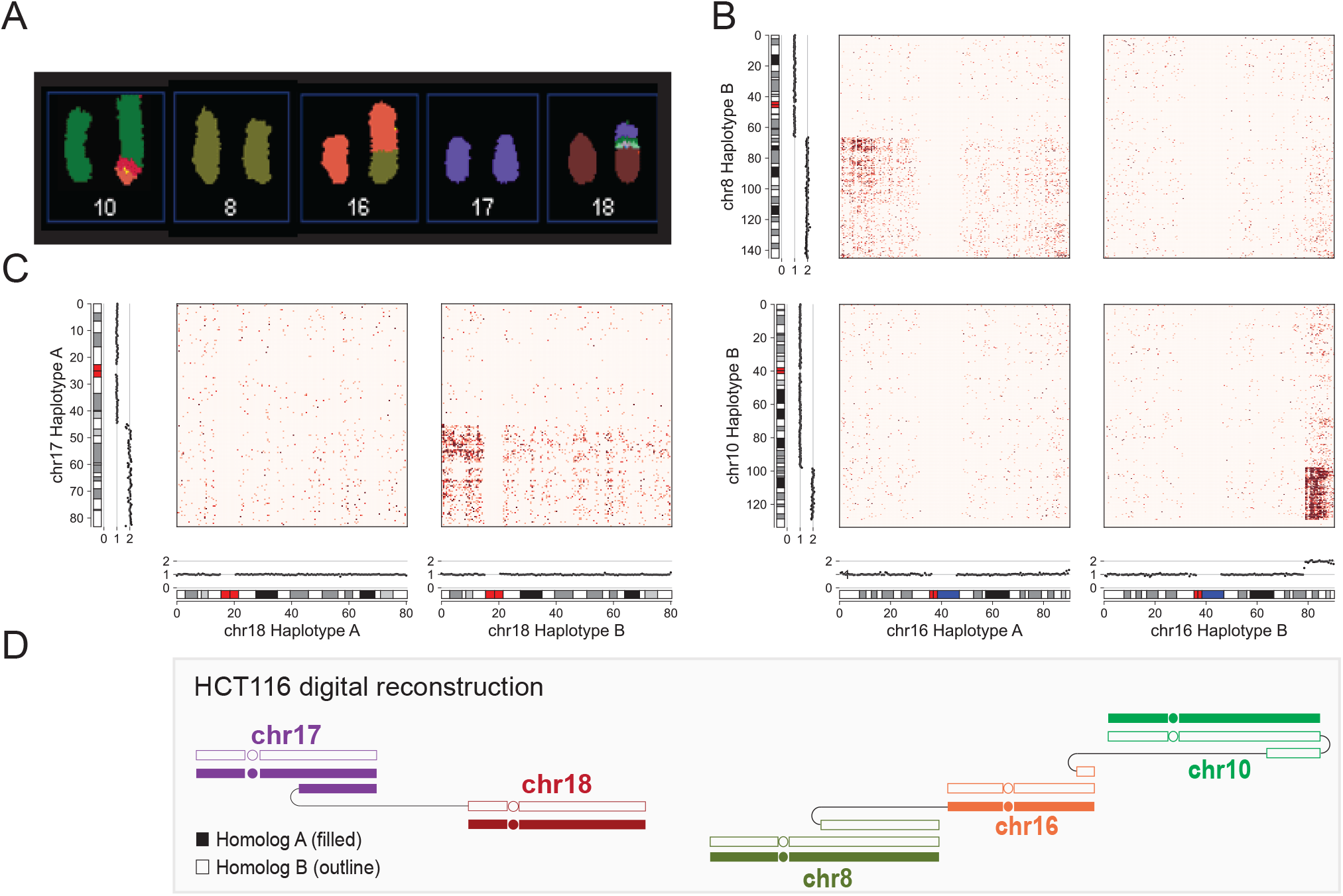
refLinker resolves the parental origin and composition of translocations in HCT116. (**A**) SKY karyotype of HCT116 rearrangements, reused from the Pawefish project with permission from Dr. Paul Edwards. (**B**) Haplotype-specific DNA copy number and inter-chromosomal Hi-C contacts between both homologs of chr16 and chr8 (top) and chr10 (bottom). DNA copy number of each homolog is in black scatterplots. Hi-C contacts are shown as heatmaps. (**C**) Haplotype-specific copy number and interchromosomal Hi-C contacts between both homologs of chr18 and chr17. Data are arranged similarly to (**B**). (**D**) Schematic drawings of HCT116 marker chromosomes. Homolog A is indicated by filled shapes and homolog B is indicated by outlines of the same color.

Based on the haplotype phase, we next calculated the copy number of every parental chromosome and identified haplotype-specific duplications on chromosome 8, 10, 16, and 17 (**Figure S5**). We then resolved the structure of translocated chromosomes based on haplotype-specific Hi-C contacts (**Figure 3B-D**). The first translocation connects a duplicated q-terminal segment of the 8B homolog (63Mb-qter) to the p-terminus of 16A, generating der(16)t(8;16) (**Figure 3A**). This is supported by the enrichment of Hi-C contacts between 16A and 8q (**Figure 3B**, top row). We inferred the breakpoint on 16A to be at the p-terminus where the Hi-C contacts show the highest density. We next identified a translocation connecting a duplicated q-terminal segment of the 16B homolog to the q-terminus of the 10B homolog gives rise to der(10)dup(10)(q?)t(10;16) (**Figure 3B**, bottom row). The duplicated 16qter segment joins a terminal duplication at the 10q terminus (**Figure S6A**), with the junction revealed to be chr10:98,953,544-chr16:79,696,003 based on bulk DNA sequencing (**Figure S6B**). Notably, the phasing of these translocations to different chr16 homologs suggests that these events occurred independently. Lastly, we determined der(18)t(17;18) arises from a translocation connecting a duplicated q-terminal segment of 17A to the p-terminus of 18B (**Figure 3C, D**).

After determining the segmental composition of translocated chromosomes in HCT116 cells, we further analyzed *de novo* chromosomal alterations [22], [24]. We calculated haplotype-specific DNA copy number from standard shotgun sequencing data of early- and late-passage clones following experimentally-induced telomere crisis [24]. Based on the copy number data, we identified concurrent deletions of the 18B p-arm and the 17A q-terminal duplication (**Figure S7**) that indicates a single arm-level deletion based on the karyotype reconstruction. We further identified examples of copy-number evolution from haplotype-specific copy-number variation between related progeny clones, which we inferred to have arisen from the formation and resolution of dicentric chromosomes (**Figure S8**). These results illustrate the power of haplotype-specific copy-number analysis for relating copy-number variation to mechanisms of chromosomal instability (**Figure S9**).

Next, we determined the segmental structure of several complex rearrangements in the near tetraploid HCC1954 genome. First, we determined the chromosomal locations of several chr17 fragments across three marker chromosomes. We determined that these fragments were all derived a single parental chromosome, consistent with ancestral chromothripsis. We then resolved the segmental structure of a marker chromosome containing a high-level (>100 copies) *ERBB2* amplicon contained within a linear homogenously staining regions [25] and capped by a 12p-terminal segment (**Figure S10**). Finally, we determined the composition of five rearranged chromosomes harboring segments from HCC1954’s two parental X chromosomes (**Figure 4**), finding that complete deletion of the Xq arm took place on the inactive X.

**Figure 4.**
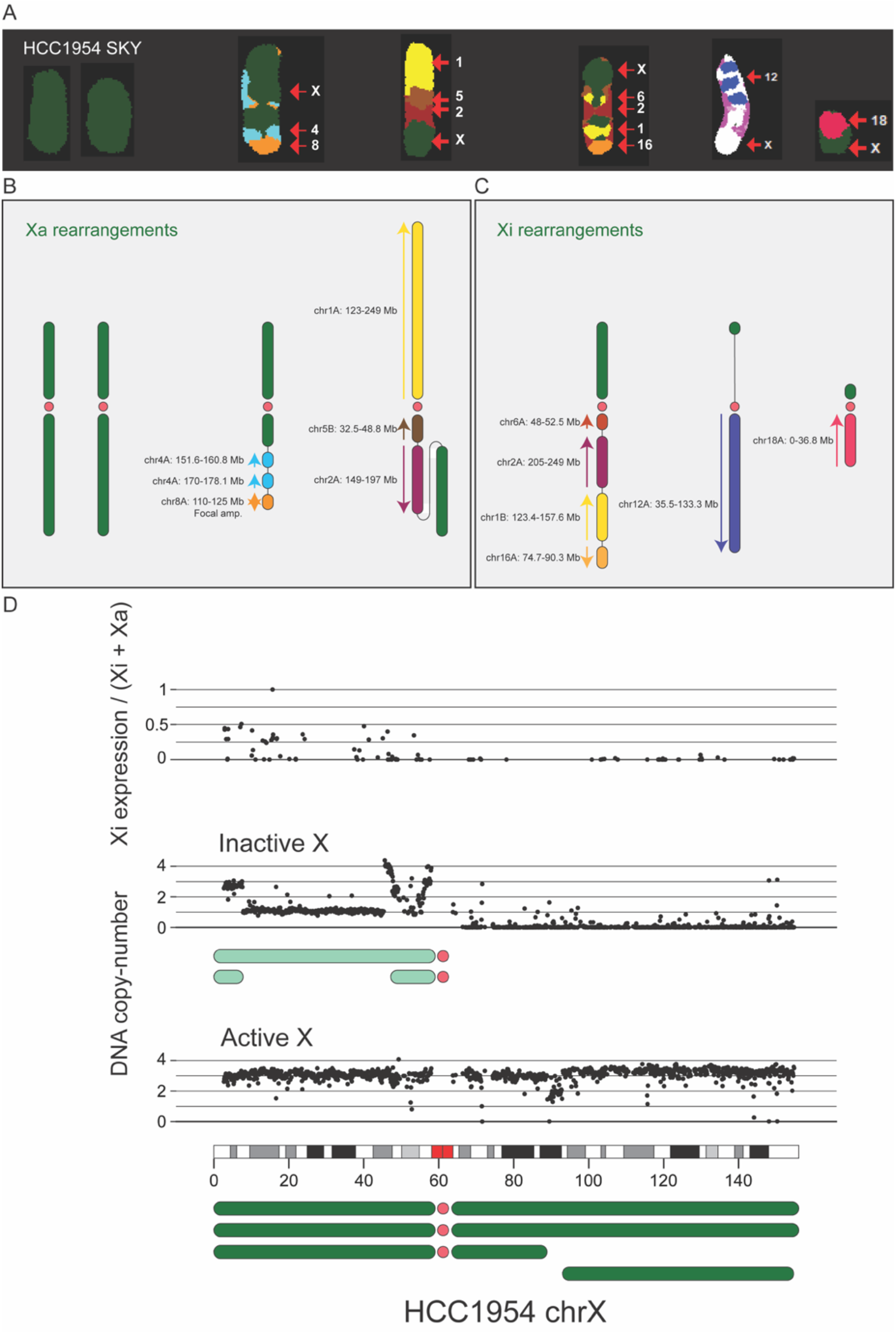
Reconstruction of X-linked rearrangements in HCC1954. (**A**) HCC1954 SKY image depicting intact and translocated X chromosomes (**B, C**) Allele-specific reconstructions of Xa- and Xi-linked rearrangements based on *refLinker* haplotypes. (**D)** Top: Relative Xi expression and Bottom: Allele-specific DNA copy-number for both chrX homologs in HCC1954.

Together, these results demonstrate a general strategy to determine the haplotype-specific segmental structure of both simple and complex rearranged chromosomes. In particular, *refLinker* enables the determination of rearrangements of both X chromosomes that have near identical sequences but distinct epigenetic and transcriptional states. We next use ths information to assess how X-chromosome rearrangements impact its transcription.

### Long-range transcriptional changes associated with X-chromosome rearrangements

In somatic cells of female eutherian mammals, dosage compensation is achieved through X-chromosome inactivation, or chromosome-wide silencing of a single X homolog [26]. X-chromosome inactivation is established during embryonic development through the upregulation of *XIST*, a long noncoding RNA located in the X-inactivation center (XIC) on the chrX q-arm. While *XIST* expression was previously considered dispensable once inactivation is established, a growing body of work suggests that *XIST* plays a critical role for the maintenance of X-chromosome silencing in somatic cells [27], [28]. In HCC1954 RNA-seq, we observed substantial transcription from segments of the inactive X chromosome contained in all three rearranged chromosomes (**Figure 4D**). As a number of X-linked genes (up to 25%) partially or completely escape X-chromosome inactivation [29], we compared the amount of Xi transcripts in HCC1954 cells to RPE-1 a genome where the inactive X chromosome is intact (**Figure 5A**). We identified two groups of genes with Xi expression. The first group exhibit biallelic expression in both HCC1954 and RPE-1, indicating that they are constitutive escapees. The second group are expressed only in HCC1954 but not RPE-1. By contrast, there is no gene showing Xi expression in RPE-1 but not in HCC1954. The higher abundance of Xi transcripts in HCC1954 cells was also observed in comparison to Xi expression in the matching lymphoblastoid cell line HCC1954BL (**Figure S11**). Together, these data indicate an increase of Xi transcription in HCC1954 cells indicating partial reversal of silencing.

**Figure 5.**
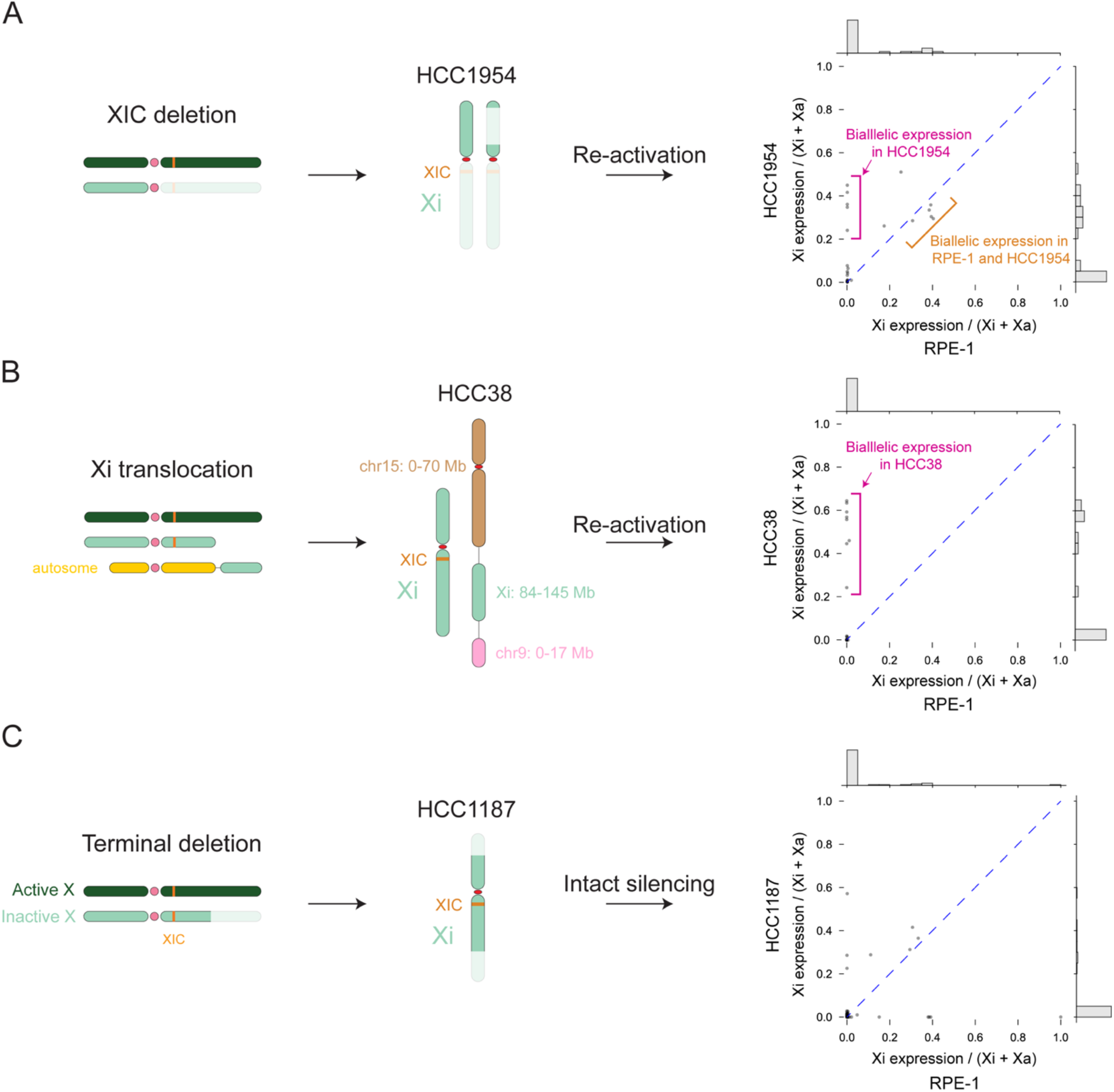
Somatic rearrangements disrupt Xi silencing in breast cancer genomes. (**A**) Left: Schematic showing deletion of the Xi q-arm, spanning the XIC. Center: Schematic of retained Xi segments in HCC1954 Right: Relative Xi expression for chrXp genes is plotted for HCC1954 (y-axis) versus RPE-1 (x-axis), a genome with an intact Xi-chromosome. Genes that escape X-chromosome inactivation in HCC1954 but not RPE-1 are indicated in magenta. Escape genes that are common to HCC1954 and RPE-1 are indicated in orange. Marginal distributions of relative Xi expression are shown for RPE-1 (horizontal histogram) and HCC1954 (vertical histogram). (**B**) Left: Schematic showing Xi q-arm translocation telomeric to the XIC. Center: Schematic of the translocated Xi segment in HCC38 Right: Relative Xi expression for chrXq genes is plotted for HCC38 (y-axis) versus RPE-1 (x-axis) Genes that escape X-chromosome inactivation in HCC38 but not RPE-1 are indicated in magenta. There are no genes in this comparison that escape inactivation in RPE-1, alone. (**C**) Left: Schematic showing terminal deletion on the Xi q-arm telomeric to the XIC. Center: Schematic of the Xi deletions in HCC1187 Right: Relative Xi expression for Xi genes is plotted for HCC1187 (y-axis) versus RPE-1 (x-axis).

We next examined Xi rearrangements and expression in the breast cancer cell line HCC38. In this genome, the p-arm of Xi is deleted, but the q-arm (including the XIC) is retained; there is also a 60Mb duplication on the telomeric side of XIC that we mapped to a derivative chromosome and flanked by segments from chr15 and chr9 (**Figure 5B, Figure S12A**). We observed a large number of transcripts from the duplicated Xq segment, none of which was expressed in RPE-1 (**Figure 5B, Figure S12B**). Notably, the higher abundance of Xi transcripts from the p-arm in comparison to the transcripts from the q-arm in RPE-1 cells is consistent with less potent silencing towards the Xp terminus [29]. Together, the abundance of transcripts from both the p-arm (in HCC1954) and the q-arm (in HCC38) of the inactive X chromosome establishes partial reversal of silencing on X chromosome fragments that are detached from XIC.

We also assessed Xi expression from fragments that retain the XIC. In HCC1187, Xi harbors terminal deletions and translocations on both the p- and q-arms, but the XIC (including *XIST*) remains intact. After measuring Xi transcription from this retained segment, we found no evidence for upregulated transcription relative to RPE-1 (**Figure 5C**).

Finally, we assessed the transcription of autosomal DNA placed in *cis* with the XIC after translocation (**Figure 6A**). In HCC38 cells, we found a derivative chromosome containing the complete Xi q-arm translocated to an ∼88Mb duplication from the chr12A q-arm, followed by a 78Mb segment from the chr4 q-arm (**Figure 6B** and **Figure S13**). To assess whether transcription of the duplicated 12q segment is suppressed by XIC in this derivative chromosome, we calculated the allelic fractions of chr12A transcripts spanning this duplication at increasing distances from the Xi junction (**Figure 6C**). We found a distance-dependent reduction of chr12A expression relative to the DNA allelic fraction (2:1 between A and B), with genes proximal to Xi showing lower transcription than genes at the distal end (**Figure 6C**). This observation suggests a direct impact on transcription arising from the XIC. We further validated this finding in HCC1187 where a truncated Xi (with the XIC intact) is flanked by a duplicated segment from the Xq arm and two duplicated segments from chr1q (150-249Mb) (**Figure 6E, Figure S14**). Analysis of the RNA/DNA copy ratio of genes on chr1 revealed a reduction of transcripts from 1q, indicating partial suppression. By contrast, the 1p arm that joins the q-arm of Xa shows a RNA/DNA ratio near one (**Figure 6F**). Together, these results suggested that Xi:autosomal translocations can cause position-dependent transcriptional suppression of the autosomal genes.

**Figure 6.**
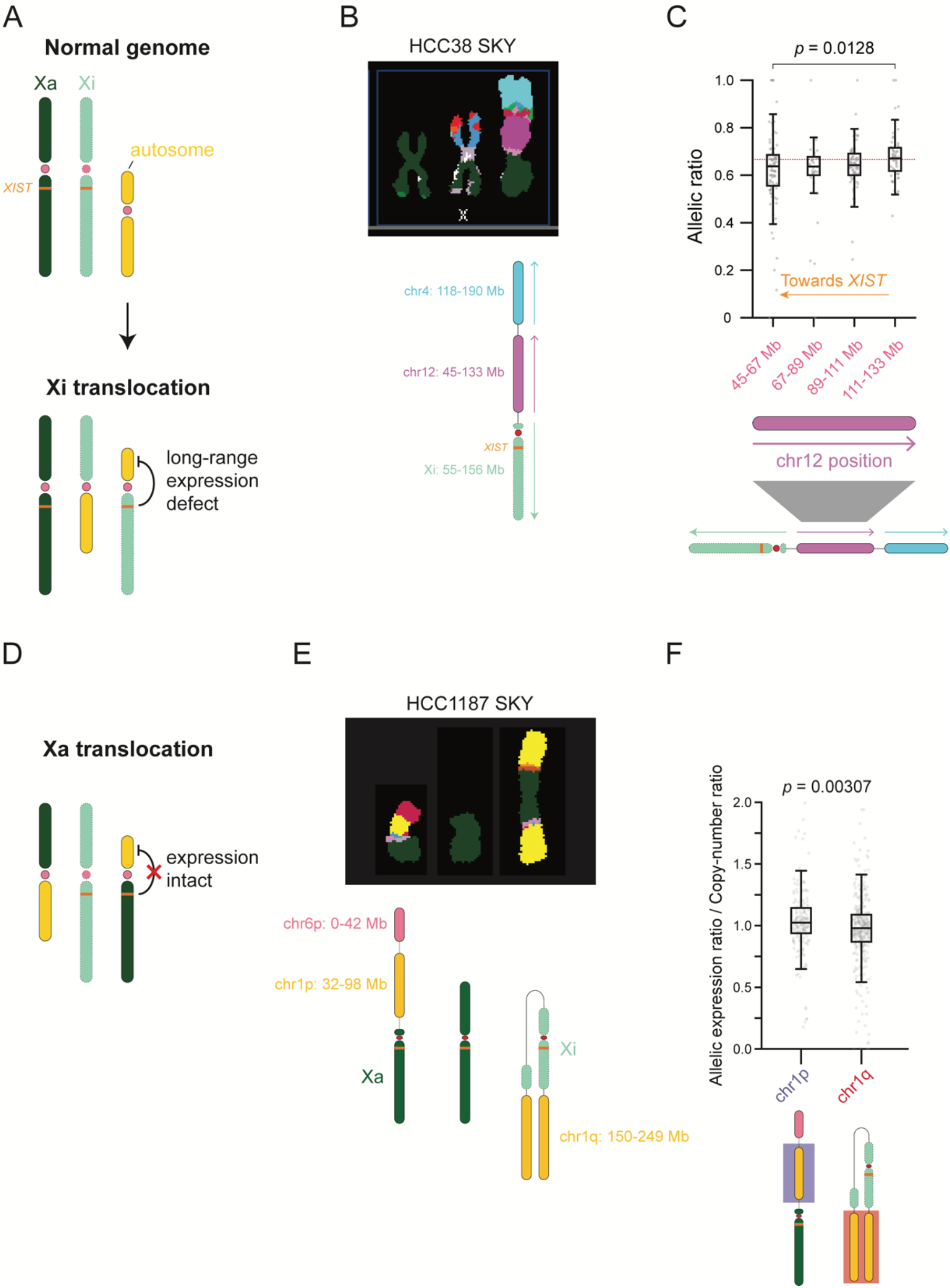
Transcriptional defects after XIST translocation. (**A**) Schematic showing X-chromosome inactivation in normal genomes (top) and after autosomal translocation to Xi (bottom) (**B**) SKY image (top) and digital reconstruction (bottom) of the marker chromosome containing Xi q-arm in HCC38. (**C**) Relative expression for 22 Mb intervals of chr12 segment translocated to Xi. Position relative to the translocated XIC is indicated by the orange arrow. (**D**) Schematic showing autosomal translocation to Xa. (**E**) SKY image (top) and digital reconstruction (bottom) of Xa- and Xi-linked rearrangements in HCC1187. (**F**) Relative expression for chr1p (left) and chr1q (right) segments translocated to Xa and Xi, respectively.

In summary, our genomic and transcriptomic analyses of Xi:autosome translocations provides direct, quantitative evidence for long-range transcription suppression of autosomal genes and partial reversal of X-chromosome silencing.

## Discussion

In a cancer genome, each rearranged chromosome consists of two or more segments from 46 parental chromosomes. To precisely quantify gene transcription from each chromosome requires two pieces of information: the haplotype phase of each parental chromosome, and the haplotype-specific segmental structure of each rearranged chromosome. Here, we present *refLinker*, a computational method that enables chromosome-scale parental haplotype inference from Hi-C data. In contrast to existing methods, *refLinker* ensures chromosome-scale haplotype accuracy from the Hi-C data of either diploid cells (e.g., germline reference) and aneuploid cancer cells.

This result demonstrates the sufficiency of bulk Hi-C sequencing for chromosome-scale haplotype inference in both diploid and structurally complex aneuploid genomes.

Based on the parental haplotype information, we further demonstrate a strategy to determine the long-range structure of rearranged chromosomes by haplotype-specific Hi-C contacts using examples from four cancer cell lines. This application can resolve the linkage between rearranged segments of 0.1-10Mb, thereby bridging the gap between sequence-level resolution from shotgun reads (10^2^-10^5^ bp) and chromosomal level resolution (10 Mb or above) from cytogenetic analysis. This strategy also reveals the parental haplotype of the rearranged segments. The determination of haplotype-resolved segmental composition of rearranged chromosomes is the first step towards end-to-end assembly of rearranged chromosomes. It also enables us to directly interrogate rearrangement-associated transcriptional changes.

The haplotype-specific reconstruction of cancer genomes is especially relevant for the analysis of rearrangements involving different X chromosomes in female cancers. As reported in a recent study from our group, both the active X and the inactive X chromosome can undergo rearrangements or translocations, but such alterations have distinct fitness effects [28]. Here, our analysis of X-chromosome rearrangement and associated transcriptional changes in three breast cancer cell lines provides definitive evidence for partial reversal of X-chromosome inactivation following XIC removal, and partial *de novo* suppression of autosomal genes translocated to the XIC. Whereas previous attempts to model these effects experimentally have led to conflicting results [13], [14], [30], we provide the first direct demonstration of both of these effects in cancer cells. Our findings identify Xi rearrangements as a potentially significant source of transcriptional dysregulation in cancer development and highlight *refLinker*’s broader utility for studying the functional consequences of cancer genome complexity.

