## Supplementary Material for "Haplotype-resolved reconstruction and functional interrogation of cancer karyotypes"

### Haplotype-resolved karyotype construction from Hi-C data using refLinker

#### This PDF file includes:

Supporting text
Figs. S1 to S14
Tables S1 to S9
Legends for Dataset S1 to S2
SI References

#### Other supporting materials for this manuscript include the following:

Datasets S1 to S2

### Supporting Information Text

### Approach

**Limitations of molecular phasing and statistical phasing.** Prior studies have demonstrated both the utility and limitations of Hi-C data for haplotype inference (1–4). Briefly, intrachromosomal Hi-C contacts can provide chromosome-scale haplotype linkage that ensures long-range phasing accuracy, but the sparsity of long-range Hi-C contacts limits the completeness of haplotype inference (1). To overcome this limitation, long reads or linked-reads are used to produce short-range haplotype blocks with dense coverage of heterozygous genotypes, which are then scaffolded by long-range Hi-C contacts (2, 3).

In addition to molecular phasing from long reads, local haplotype linkage between common variants can be inferred from reference haplotypes in a genotyped population by statistical phasing (5, 6). However, haplotype inference from statistical phasing is prone to long-range switching errors due to recombination. To evaluate the long-range accuracy of statistical phasing, we compared results from two well-established methods, EAGLE2 (6) and SHAPEIT2 (5) to truth haplotype data in the RPE-1 cell line that we determined previously from linked-reads and Hi-C (3). The EAGLE2 and SHAPEIT2 statistical haplotypes were generated using reference haplotype data from the 1000 Genomes Project Phase 3 (7) on heterozygous variants in RPE-1. Results generated by both tools show highly accurate linkage between variants within 1 kb ($> 95\%$), with EAGLE2 showing better overall accuracy than SHAPEIT2 (Figure S1A). However, the accuracy drops significantly for variants that are further apart ($70 - 80\%$ at 10 kb). Moreover, the accumulation of switching errors over many variants prevents long-range haplotype inference over intervals $> 1$ Mb (Figure S1B).

We reasoned that long-range switching errors in statistical phasing haplotypes may be corrected with Hi-C linkage to achieve chromosome-scale accuracy. The feasibility of this strategy is demonstrated by a recent method, IntegratedPhasing, which samples statistical phasing linkage from the SHAPEIT2 haplotype graph for integration with Hi-C reads, achieving accurate long-range haplotype inference in the diploid genome, NA12878 (4). However, because this approach relies on the assumption of uniform sequencing coverage between homologs and across the genome, it remained unclear whether the same strategy could be applied to aneuploid cancers. To overcome this limitation, we developed `refLinker` (https://github.com/gbrunette/refLinker), a new approach for complete haplotype inference from statistical phasing and Hi-C that is directly applicable to aneuploid genomes. We further perform a systematic evaluation of the long-range phasing accuracy of IntegratedPhasing and `refLinker` using Hi-C data of both diploid genomes and aneuploid cancer genomes with deletions, duplications, and higher-level copy-number alterations.

**A general formalism of integrative haplotype inference.** As described in our previous study (3), we use a binary representation for heterozygous genotypes and their corresponding haplotype phase. Briefly, reference and alternate genotypes at a heterozygous position $i$ are represented as $\sigma_i = +1$ or $-1$, respectively and the chromosomal haplotype is represented as the vector

$$\mathbf{S} = (s_1, s_2, \dots s_N) \quad s_i = \pm 1. \tag{1}$$

Molecular linkage from Hi-C reads spanning two or more heterozygous sites is similarly represented as

$$\mathbf{\Sigma} = \{\sigma_i | 1 \leq i \leq N\} \quad \sigma_i = \pm 1. \tag{2}$$

This formalism reduces haplotype inference to finding $\mathbf{S}$ that maximizes

$$\sum_{k=1}^{M} \sum_{1 \leq i,j \leq N} \sigma_{i,k}\sigma_{j,k} s_i s_j, \tag{3}$$

where the first sum is carried over all molecular links and the second over all variant sites. The maximization is equivalent to finding the haplotype phase with the lowest number of clashes, i.e.

$$s_i s_j \sigma_{i,k}\sigma_{j,k} = -1.$$

We further introduce the aggregate molecular linkage between two sites $i$ and $j$ as

$$M_{ij} = \chi_{ij} \sum_{k=1}^{m} \sigma_{i,k}\sigma_{j,k}, \tag{4}$$

with a weighting factor

$$\chi_{ij} = \frac{1}{m} \left| \sum_{k=1}^{m} \sigma_{i,k}\sigma_{j,k} \right| \in [0, 1] \tag{5}$$

that measures the concordance of links between two variant sites; $\chi_{ij}$ serves to penalize linkage evidence from false variant sites with links to both parental haplotypes.

The optimal haplotype solution based on Hi-C links should maximize:

$$E(\mathbf{S}) = \frac{1}{2} \sum_{ij} M_{ij} s_i s_j. \tag{6}$$

Where, for $\chi_{ij} \approx 1$, we have

$$E(\mathbf{S}) \approx (\#\ \text{links consistent with}\ \mathbf{S}) - (\#\ \text{links inconsistent with}\ \mathbf{S})\,. \quad [7]$$

Let $\mathbf{G} = (g_1, g_2, \cdots g_N), g_i \in \{-1, 1\}$ be the statistical phased haplotype. We can introduce statistical haplotype linkage into Eq. (6) as

$$E(\mathbf{S}) = \frac{1}{2} \sum_{1 \le i,j \le N} (M_{ij} + \rho_{ij} g_i g_j)\, s_i s_j. \quad [8]$$

Here $\rho_{ij}$ is the "strength" of haplotype linkage from statistical phasing for sites $i$ and $j$. Based on the decay of statistical phasing accuracy shown in Figure S1, we assume an exponential function of $\rho_{ij}$ reflecting the accumulation of switching errors:

$$\rho_{ij} = e^{-d_{ij}/c}, \quad [9]$$

where $d_{ij}$ is the 1D genomic distance between variant sites $i$ and $j$ and the constant $c$ measures the average genetic distance of recombination. Because statistical phasing mainly relies on linkage disequilibrium between adjacent variants, we only incorporate linkage between adjacent variants. This leads to the following function to be maximized:

$$\begin{aligned} E(\mathbf{S}) &= \frac{1}{2} \left( \sum_{1 \le i,j \le N} M_{ij} s_i s_j + \sum_{1 \le i < N} \rho_i g_i g_{i+1} s_i s_{i+1} \right) \\ &= \frac{1}{2} \sum_{1 \le i,j \le N} M'_{ij} s_i s_j. \end{aligned} \quad [10]$$

Here $\rho_i = \rho_{i,i+1} = e^{-d_{i,i+1}/c}$.

**Using Hi-C reads to correct long-range switching errors.** For haplotype determination using Hi-C and long reads, we previously maximized Eq. (6) by iteratively considering two perturbations to a random initial guess $\mathbf{S}_0$: a local spin flip

$$(s_1, s_2, \cdots, s_i, \cdots, s_N) \to (s_1, s_2, \cdots, -s_i, \cdots, s_N),$$

and a global switch

$$(s_1, s_2, \cdots, s_i, s_{i+1}, \cdots, s_N) \to (s_1, s_2, \cdots, s_i, -s_{i+1}, \cdots, -s_N).$$

To correct switching errors of large haplotype blocks, we consider "block flips" defined as

$$(s_1, s_2, \cdots s_k, s_{k+1} \cdots s_l, s_{l+1}, \cdots s_N) \to$$

$$(s_1, s_2, \cdots s_k, -s_{k+1} \cdots - s_l, s_{l+1}, \cdots s_N)$$

A block flip is equivalent to two global switches at site $k$ and site $l$ in a single step (Figure S1C). Starting with $\mathbf{S_0} = \mathbf{G}$, we find $\hat{\mathbf{S}}$ that maximizes Eq. (10) by applying block flips.

For a block flip between site $k+1$ and $l$, we have

$$\Delta E_{k|k+1,l|l+1} = - \sum_{k < i \le l} s_i \left( \sum_{j > l} M'_{mj} s_j + \sum_{j \le k} M'_{mj} s_j \right). \quad [11]$$

This can be calculated using a recursive relationship

$$\begin{aligned} \Delta E_{k|k+1,l|l+1} - \Delta E_{k-1|k,l-1|l} = {} & s_k \left( \sum_{i>l} + \sum_{i<k} - \sum_{k<i<l} \right) M'_{ik} s_i \\ & - s_l \left( \sum_{i>l} + \sum_{i<k} - \sum_{k<i<l} \right) M'_{il} s_i. \end{aligned} \quad [12]$$

Note that $\Delta E_{0|1,l|l+1}$ is the block switching penalty $\Delta E_{l|l+1}$ given by

$$\Delta E_{l|l+1} = - \sum_{i \le l} \sum_{j > l} M'_{ij} s_i s_j. \quad [13]$$

and can be calculated using

$$\Delta E_{l|l+1} - \Delta E_{l-1|l} = s_l \left( \sum_{j>l} M'_{lj} s_j - \sum_{i<l} M'_{il} s_i \right). \quad [14]$$

We use this recursion to calculate block flipping penalties. For each step of haplotype refinement, block flipping penalties are calculated at every variant position for block sizes ranging from half the total number of variants on a chromosome ($N/2$) down to a single variant. Specifically, we calculate large-scale flipping penalties for block sizes in the sequence $N/2, N/4, \ldots, \lceil N/2^{i+1} \rceil > 2000$. Finer-scale flipping penalties are calculated for block sizes on the interval $[2000, 200]$ with a 20 variant decrement and on the interval $[200, 1]$ with a 1 variant decrement.

This results in a grid of block flipping penalties for every position on the chromosome, for every block size described above. We then simultaneously flip the haplotype assignment for disjoint blocks with the most favorable switching penalties satisfying $\Delta E \geq 10.0$. This process is repeated until no remaining favorable block flips are found.

The resulting solution only includes phased genotypes at variant sites with Hi-C linkage (see Table S2,"linked by Hi-C"). For variants not having Hi-C links, we determined their haplotype phase based on their statistical phasing linkage to the nearest Hi-C phased variant (up to 5kb, see Table S2 "EAGLE2 recovery"). The combination of Hi-C phasing and local haplotype recovery produces a chromosomal haplotype with phased genotypes at $> 99\%$ common variant sites ($\sim 1$ variant per 1.5 kb).

### Results

**Benchmarking IntegratedPhasing in diploid genomes.** We compared `refLinker`'s phasing accuracy to IntegratedPhasing (4). IntegratedPhasing and `refLinker` differ in both (1) their approaches for integrating haplotype linkage from statistical phasing; and (2) their quantification of Hi-C linkage. For diploid phasing applications, the first point is most relevant: because IntegratedPhasing uses the SHAPEIT2 haplotype graph to extract high-confidence statistical phasing linkage, it is less flexible to the incorporation of improved statistical phasing algorithms, such as EAGLE2 (Figure S1A). For comparison, we ran IntegratedPhasing on Hi-C data from the RPE-1 genome (Table S1). After running on complete Hi-C data from RPE-1, IntegratedPhasing achieved 98.7% accuracy. However, IntegratedPhasing's lower overall accuracy on downsampled Hi-C data (95.9%, including seven chromosomes with $> 5\%$ phasing errors after $0.333\times$ downsampling; Table S8) indicates that IntegratedPhasing requires more long-range Hi-C contacts than refLinker for robust, long-range haplotype determination.

Next, we evaluated each method's robustness across large variant deserts or gaps in the human genome, including centromeres. Having established that refLinker requires $\sim 10$ million long-range Hi-C contacts for robust haplotype inference, we ran both refLinker and IntegratedPhasing on $0.333\times$ downsampled Hi-C data from RPE-1. We then examined switching errors across HSat3 on chr9, a 27.6 Mb repetitive array that leads to spurious errors in linking p- and q-arm haplotypes (1). For 20 independent trials of $0.333\times$ Hi-C downsampling, three runs from IntegratedPhasing returned arm-level switching errors (Figure S2A, left), whereas refLinker returned none (Figure S2A, right). The accuracy of refLinker haplotypes ranged from $97.5 - 98.6\%$ (Figure S2B).

### Methods

**refLinker implementation.** We implemented the `C++` package `refLinker` for whole-chromosome haplotype determination from Hi-C data and statistical phasing (https://github.com/gbrunette/refLinker). First, Hi-C linkage between genetic variants is determined with the command "`reflinker extract`". As described in Ref (3), "`reflinker extract`" takes aligned Hi-C reads and a list of variant genotypes to generate a map from variant genotypes to the Hi-C sequencing reads that cover them. "`reflinker solve`" then takes the resulting map and a statistical phasing solution and minimizes $\Delta E(\mathbf{S})$. Given aligned Hi-C reads and statistical phasing solution, `refLinker` phasing runs with the commands:

```
# Generating the reads to variant map: {HiC variant map}.dat
reflinker extract -v {statistical phasing haplotype}.vcf.gz -i {HiC reads}.bam -e hic;
# Generating whole-chromosome haplotypes
reflinker pop -v {statistical phasing haplotype}.vcf.gz -g {HiC variant map}.dat -e -10.0 -p 0.999;
```

Where the flag `-e` sets the $\Delta E(\mathbf{S})$ cutoff for haplotype refinement (default value $-10.0$) and `-p` sets a pruning parameter, removing outlier variants with unusually high Hi-C linkage. Running on the default "`-p 0.999`," `reflinker solve` prunes outlier variants that exceed the $99.9^{\text{th}}$ percentile in Hi-C linkage. This mitigates the effect of Hi-C coverage artifacts on haplotype determination.

"`reflinker pop`" generates a haplotype scaffold consisting of phased variants linked by Hi-C. We provide the python script `eagle2_recovery.py` to recover unlinked, common germline variants. `eagle2_recovery.py` calculates the haplotype linkage between each unlinked variant in the `EAGLE2` haplotype and the closest linked variant within 5 kb. Given an unlinked variant, $i$, and the nearest linked variant, $j$, we recover the missing haplotype assignment as: $s_i = g_i g_j s_j$.

**Sequence alignment and post-processing.** Hi-C and shotgun sequencing data were aligned using `bwa mem` with default parameters (8). As in Ref (3), read-pair configuration was used to assign primary alignment positions to sequencing reads with supplementary or secondary alignments. PCR duplicates were tagged using Picard `MarkDuplicates`. Aligned bams were downsampled with the Picard program `DownsampleSam`.

**Variant calling and filtering.** For RPE-1 and NA12878, genotypes were used from Ref (3). For HCC1954BL and HCT116 GATK `HaplotypeCaller` was run in discovery mode ("`--genotyping-mode DISCOVERY`") to genotype variants from shotgun sequencing data. We ran `HaplotypeCaller` with the following read filters:

```
--read-filter PairedReadFilter \
```

```
--read-filter MateOnSameContigOrNoMappedMateReadFilter \
--read-filter FragmentLengthReadFilter --max-fragment-length 1000 \
--read-filter MateDifferentStrandReadFilter \
--read-filter MappingQualityReadFilter --minimum-mapping-quality 30 \
--read-filter OverclippedReadFilter --filter-too-short 25 \
--read-filter GoodCigarReadFilter --read-filter AmbiguousBaseReadFilter
```

For HCT116 genotyping was performed jointly on shotgun sequencing samples SRR6312252, SRR6312255, SRR6312256, SRR6312257, SRR6312258, from (9). For HCC1954BL genotypes were determined from shotgun sequencing generated in this study.

**Evaluating `refLinker` performance.** We ran `refLinker` on three diploid human genomes with ground truth haplotype information to benchmark the accuracy and completeness of haplotype inference from Hi-C data (see Table S1 for Hi-C data sources). For NA12878, truth haplotypes were determined using parental genomes in the Genome In A Bottle release (10); for RPE-1 cells, truth haplotypes were determined from single-cell sequencing and independently validated by computational inference from linked reads and Hi-C reads with mLinker (3); for HCC1954BL, truth haplotypes were determined from linked-reads and Hi-C data with mLinker and validated by allelic imbalance in the matching breast cancer genome HCC1954. Phasing accuracy was evaluated both for variants with direct haplotype linkage from Hi-C reads (Table S2: “linked by Hi-C”) and including variants with only statistical haplotype linkage to Hi-C phased variants (Table S2: “recovered”). In all cases, refLinker was able to resolve the haplotype phase of common germline variants at $> 99\%$ accuracy (Table S3 S4 S5).

**Benchmarking haplotype phasing algorithms.** The statistical phasing algorithms `EAGLE2` (11) and `SHAPEIT2` (12) were run with default parameters on reference panel data from 1000 Genomes Project Phase 3 (13). For `EAGLE2`, we used the GRCh38 liftover. VCF outputs were generated from `SHAPEIT2` with the command:

```
shapeit -convert --input-haps {sample}.phased --output-vcf {sample}.phased.vcf
```

We ran the `IntegratedPhasing` pipeline (14) on RPE-1 Hi-C and genotype data. Hi-C fragments were extracted using the HapCUT2 `ExtractHAIRS` program with the Hi-C option (“`--hic 1`”) (15). RPE-1 variants were lifted over to GRCh37 using Picard `LiftoverVCF` to run `SHAPEIT2`. Hi-C fragments and “pseudo-reads” generated in the `IntegratedPhasing` pipeline were then concatenated to run `HapCUT2` with the Hi-C option (“`--hic 1`”).

For `refLinker` benchmarking, Hi-C fragments were extracted using the `mlinker extract` program (16). `refLinker` was run with $\Delta E(\mathbf{S}) \leq -10.0$ cutoff for full Hi-C datasets (i.e. “`-e -10.0`”) and $-5.0$ cutoff for down-sampled Hi-C data (i.e. “`-e -5.0`”).

**Metaphase Preparation and Spectral Karyotyping (SKY).** Metaphases chromosomes were prepared by incubating 100ng/ml Colcemid (Roche, Brighton, MA) for two hours followed by mitotic shake off. Mitotic cells were then treated with a hypotonic solution (0.075M KCl) for 20 minutes at 37° C. Subsequently, the cells were centrifugated, supernatant extracted, fixed and rinsed three times with methanol/acetic acid (3:1) before being fixed onto a slide within a humidity-controlled Thermotron (Thermotron, Holland MI).

SKY probes using a combination of five different fluorochromes were prepared in-house as previously described Ref. (17). Chromosome labeling was performed by directly incorporating Dy-505 (Dyomics, Jena, Germany), Spectrum Orange-dUTP (Abbott Molecular, Des Plaines, IL) Dy-590, Biotin-16-dUTP (Sigma-Aldrich, Inc, St. Louis, MO), and Digoxigenin-11-dUTP (Sigma-Aldrich) in a secondary PCR reaction followed by detection using Avidin Cy5 (Rockland Immunochemicals, Limerick, PA), Mouse Anti-Digoxin antibody (Sigma-Aldrich) and a goat antimouse-antibody conjugated to Cy5.5 (Rockland Immunochemicals). Slides were mounted and counterstained with DAPI (Vector Laboratories, Newark, CA). Hybridization occurred over a period of 2 days at 37°C. Detailed protocol A minimum of 15 metaphases were imaged and karyotyped using ASI GenASIS 8.2.2 software on the Olympus BX63 microscope (Evident, Tokyo, Japan) equipped with a Spectral Cube (Applied Spectral Imaging, Carlsbad, CA).

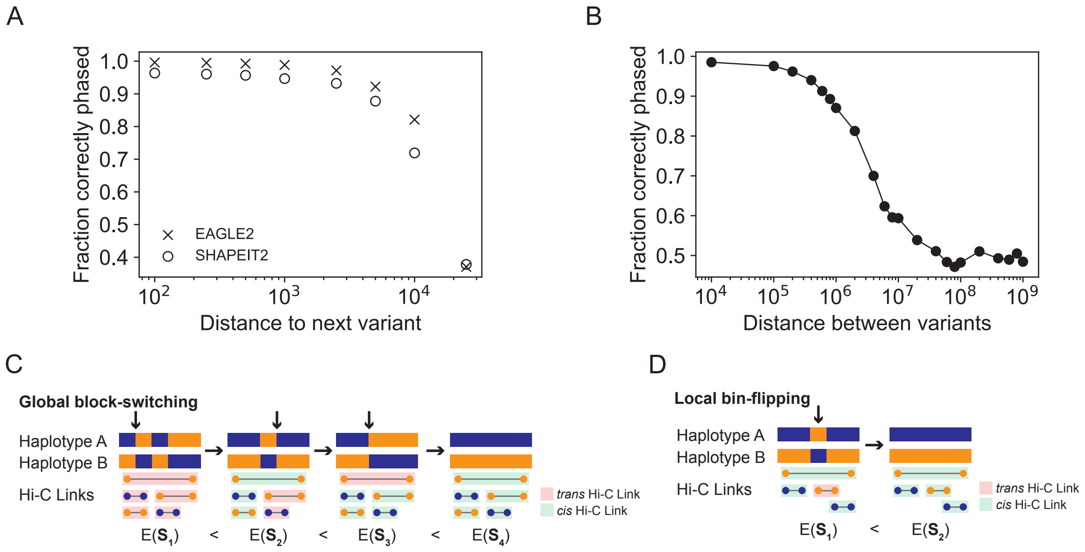


**Fig. S1.** Determination of whole-chromosome haplotypes from statistical phasing and Hi-C. (A) Fraction of correct haplotype linkage is plotted for adjacent variants on RPE-1 chr2 at different genomic distances for the statistical phasing algorithms EAGLE2 (×) and SHAPEIT2 (○). (B) Fraction of correct haplotype linkage derived by EAGLE2 for all variants on RPE-1 chr2. For (A) and (B), phasing accuracy was assessed using truth data from experimental phasing and only for single-nucleotide polymorphisms. (C) Schematic of the strategy to correct long-range switching errors by minimizing apparent trans Hi-C contacts. The starting point is the haplotype from statistical phasing (left), with blue and orange representing the true parental haplotype (right). Switching errors are indicated by transitions between blue and orange. Hi-C links are shown below as pairs of blue or orange circles. Hi-C links indicating cis contacts (A-A or B-B) are colored green and trans contacts (A-B) are colored red. refLinker iteratively searches for switching blocks (arrows) to minimize the number of apparent trans Hi-C links.

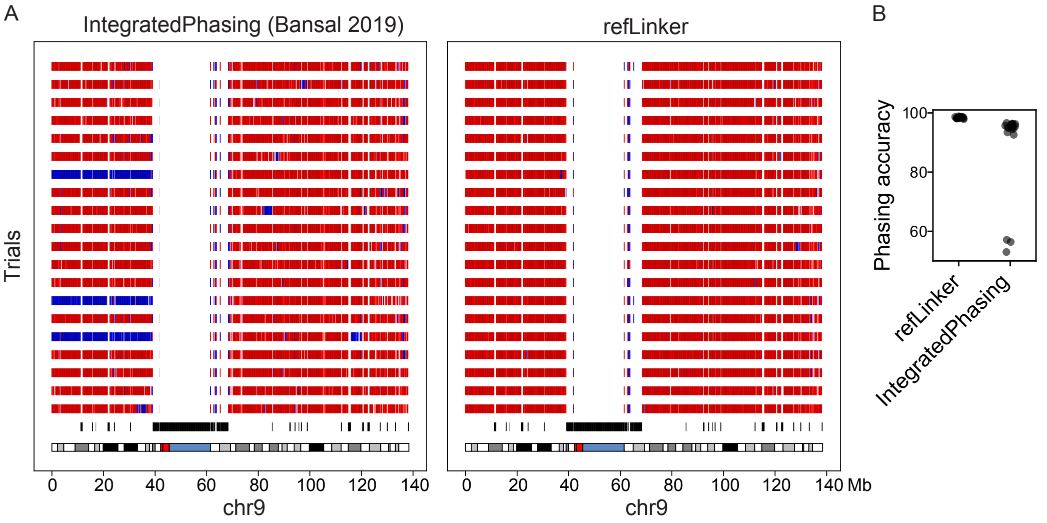


**Fig. S2.** Haplotype phasing over satellite repeats. (A) `refLinker` and `IntegratedPhasing` were run on 20 trials of $0.333\times$ downsampling of RPE-1 Hi-C data. For both, variant density is shown on the bottom track, where black bins indicate variant-poor regions, including the chr9 centromere and 27.6 Mb contiguous satellite array. Red-to-blue switch-points indicate long-range phasing errors. (B) Whole-chromosome phasing accuracy is plotted for 20 rounds of downsampling for `refLinker` and `IntegratedPhasing`.

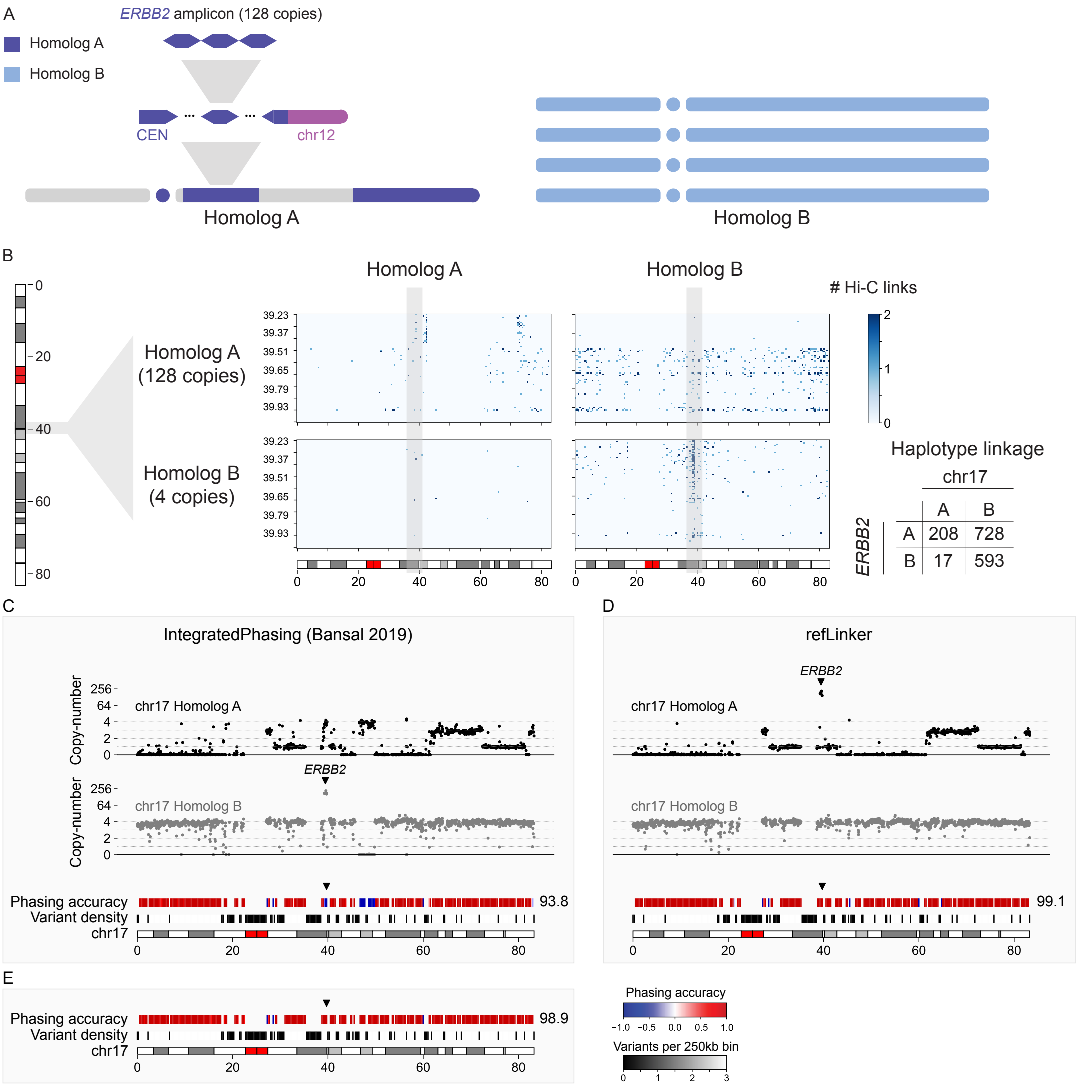


**Fig. S3.** (A) Schematic showing HCC1954 chr17 A and B homologs (dark blue and light blue, respectively). Haplotype resolved DNA copy-number reveals Homolog A is highly rearranged with p-arm deletion, large deletions on the q-arm and and *ERBB2* focal amplification (left). In contrast, Homolog A is present in four intact copies (right). (B) Haplotype-specific Hi-C contact map between the *ERBB2* locus and the rest of chr17, where the total number of Hi-C contacts are: 17B-*ERBB2*↔17A (728), 17A-*ERBB2*↔17A (593), 17B-*ERBB2*↔17B (208) and 17A-*ERBB2*↔17B (17). (C) IntegratedPhasing was run on HCC1954 Hi-C data. Allelic copy-number is plotted for chr17 Homolog A (top row, gray) and Homolog B (second row, black). Phasing accuracy and variant density is plotted for 250 kb bins (bottom two tracks). Whole-chromosome IntegratedPhasing accuracy for chr17 was $93.8\%$. Focal amplification containing *ERBB2* was assigned to Homolog A (black arrow). (D) refLinker was run on HCC1954 Hi-C data and initialized with EAGLE2 haplotypes for HCC1954BL. Allelic copy-number is plotted for chr17 Homolog A (top row, gray) and Homolog B (second row, black). Phasing accuracy and variant density is plotted for 250 kb bins (bottom two tracks). Whole-chromosome refLinker phasing accuracy for chr17 was $99.1\%$. Focal amplification containing *ERBB2* was assigned to Homolog B (black arrow). (E) refLinker was run on HCC1954 Hi-C data and initialized with IntegratedPhasing haplotype from (G). Phasing accuracy (top track)and variant density (bottom track) is plotted for 250 kb bins. Whole-chromosome refLinker phasing accuracy for chr17 was $98.9\%$. Phasing accuracy at the *ERBB2* locus is indicated by the black arrow.

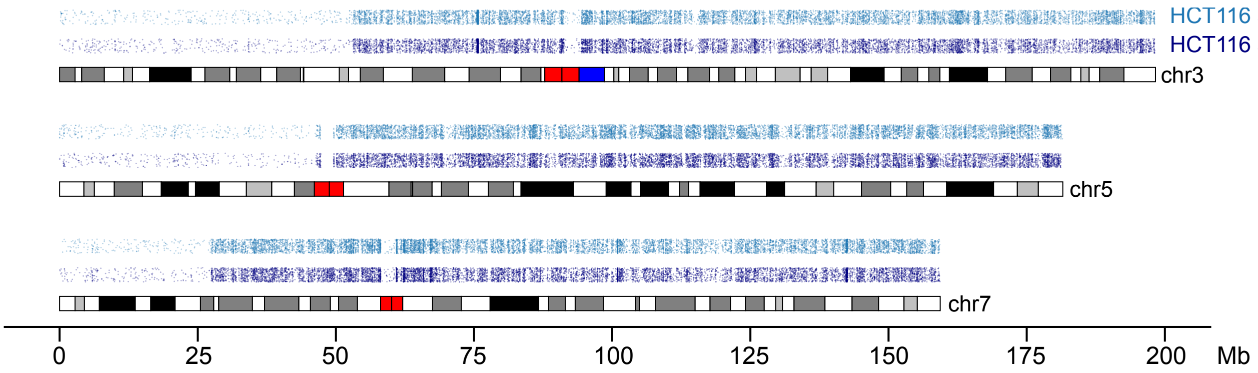


**Fig. S4.** HCT116 loss-of-heterozygosity. Heterozygous sites are plotted versus position for chr3, chr5, and chr7. Heterozygous genotypes were called from whole-genome sequencing of HCT116 from two different studies: Sample 1: SRR6312252 (9) Sample 2: SRR21384125 (18). Loss-of-heterozygosity is evident from the absence of heterozygous variants at the p-terminus of all three chromosomes.

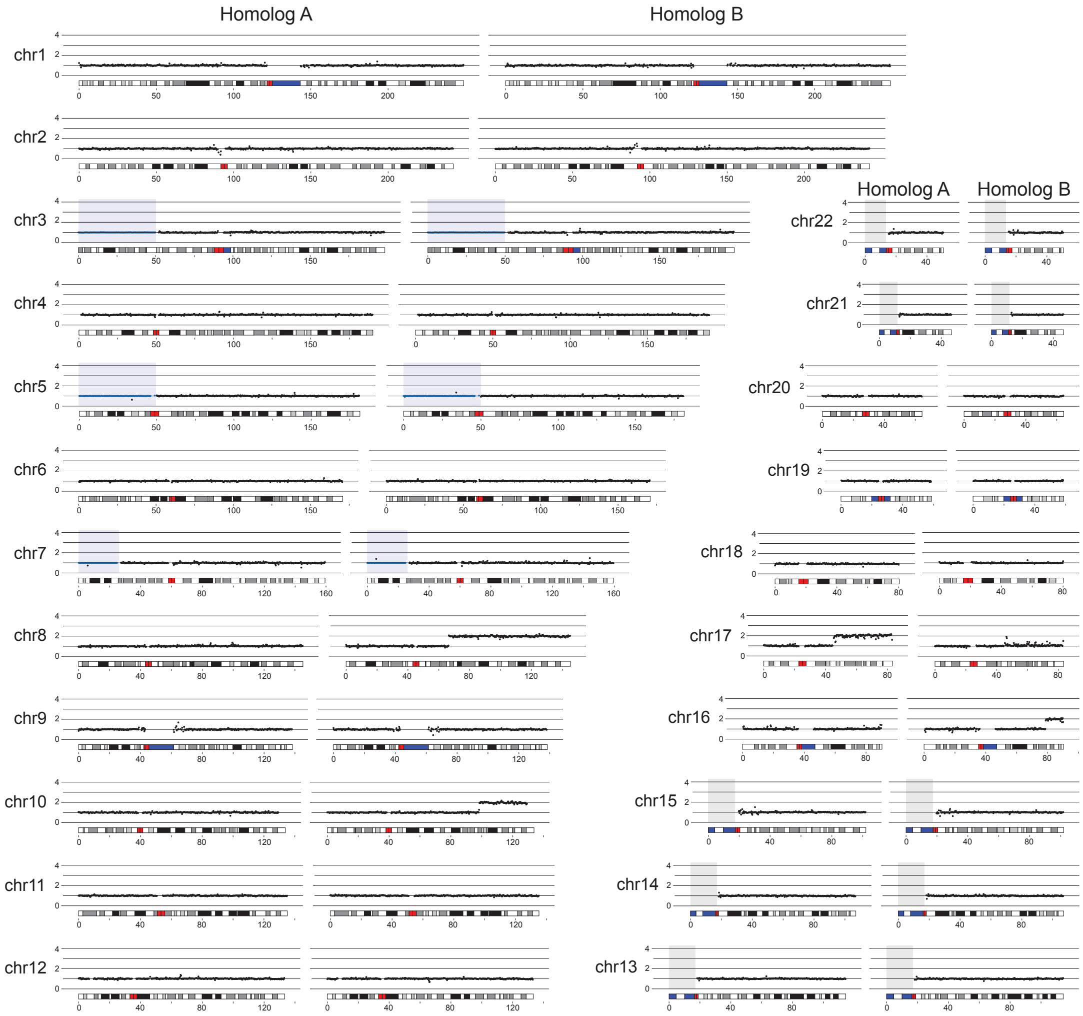


**Fig. S5.** HCT116 allelic copy-number is plotted for every autosome in HCT116. For each chromosome, Homolog A DNA copy-number is plotted on the left and Homolog B on the right. Blue shading indicates regions of copy-number neutral loss-of-heterozygosity on chromosomes 3, 5, and 7, where $1/2$ of the total copy-number is plotted. Gray shading indicates the p-arms of acrocentric chromosomes, consisting of unmappable rDNA repeats.

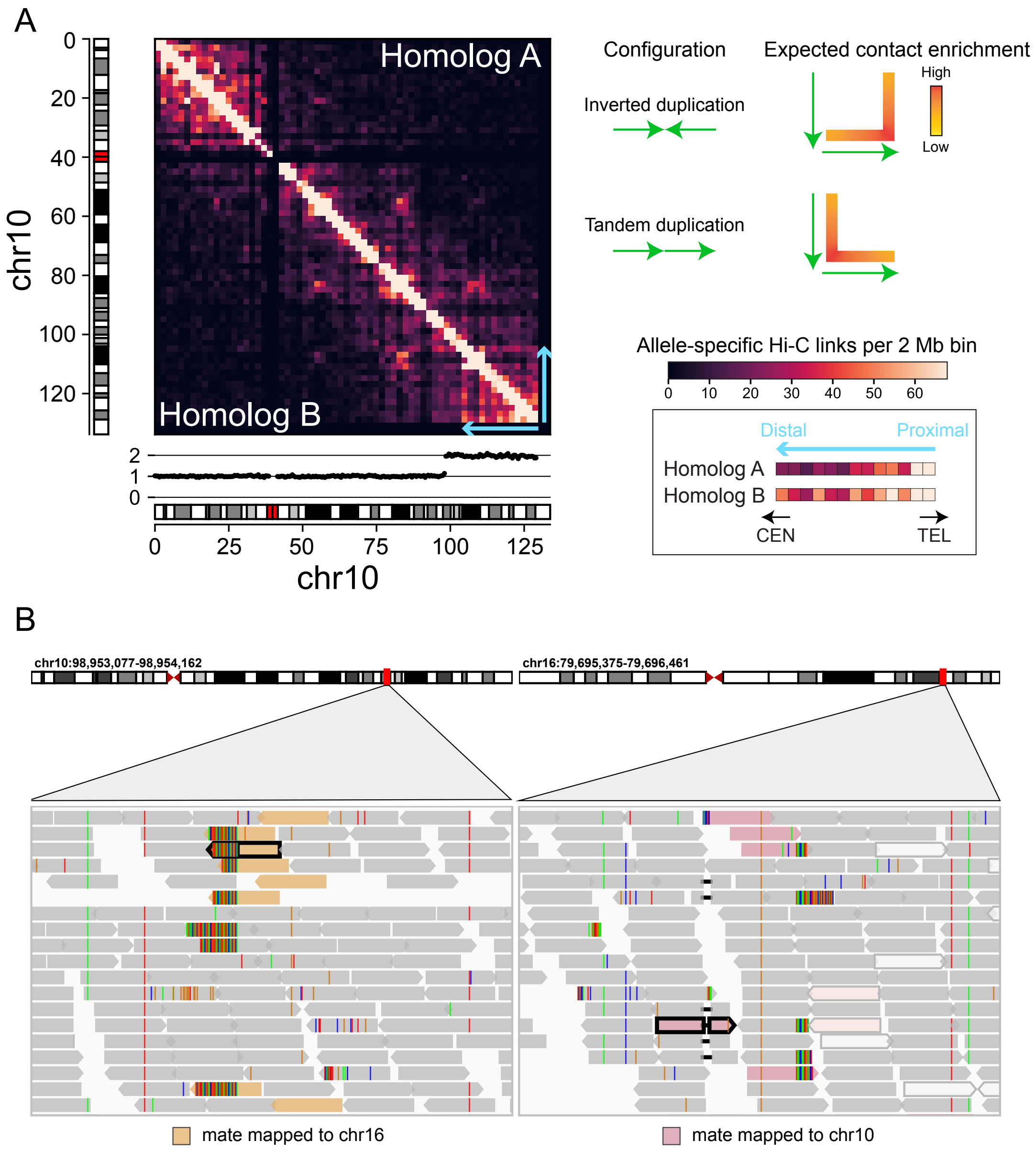


**Fig. S6.** Hi-C maps reveal HCT116 intra-chromosomal rearrangements. (A) Left: HCT116 chr10 homolog-specific intra-molecular Hi-C contacts after copy-number normalization (2 Mb bins). Labels indicate the Homolog A Hi-C linkage (upper triangle) and Homolog B Hi-C linkage (lower triangle). Cyan arrows indicate the location of chr10 duplication and orientation of the representative rows shown on the lower right. Right: (Top) Expected Hi-C contact enrichment is depicted to two possible orientations of segmental duplication, inverted duplication or fold-back (top) and tandem duplication (bottom). (Bottom) Genomic intervals are plotted for the telomeric end of HCT116 duplication for Homolog A (wild-type) and Homolog B (duplicated) after normalization for DNA copy-number. Homolog B Hi-C contact frequencies are consistent with the inverted duplication orientation. (B) Paired-end sequencing of HCT116 reveals chr10:98,953,544-chr16:79,696,003 breakpoint. Left: Sequencing reads aligned to the interval chr10: 98,953,077-98954,162. Discordant reads with mates mapped to chr16 are indicated in beige. Right: Sequencing reads aligned to the interval chr16: 79,695,375-79,696,461. Discordant reads with mates mapped to chr10 are. indicated in pink. A representative read pair is indicated by black outline.

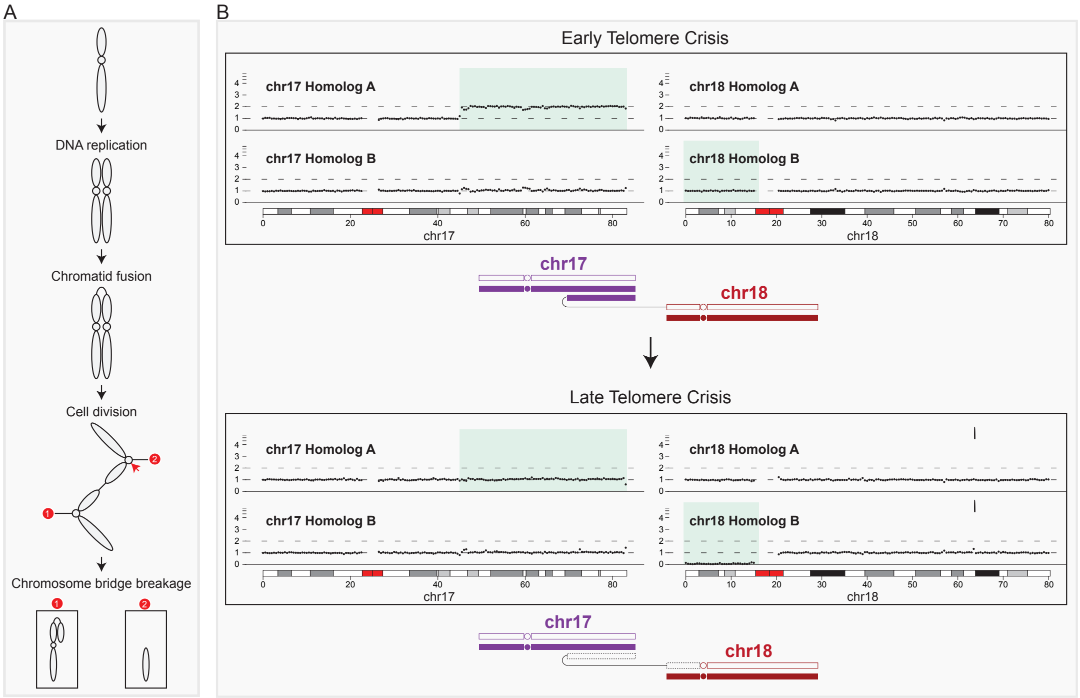


**Fig. S7.** (A) A drawn schematic showing reciprocal arm-level gain and loss resulting from a single round of chromatid-type breakage-fusion-bridge cycle. (B) chr17A-chr18B linkage revealed in HCT116 evolution following telomere crisis. Homolog specific copy-number was calculated from whole-genome sequencing of HCT116 clones at two timepoints after escape from telomere crisis. Top: Whole-genome sequencing of SRR18252999, HCT116 early telomere crisis clone, reveals chr17 and chr18 copy-number consistent with HCT116 karyotype. Drawn schematics indicate the two homologs of chr17 and molecular linkage between the chr17A duplicated segment with chr18B. Bottom: Whole-genome sequencing of the same clone at a later timepoint (SRR18252998) reveals a single-copy loss of the chr17A duplicated segment (top plot) linked to loss of the chr18B q-arm (bottom plot). Drawn schematics indicating arm-level loss on the chr17A-chr18B derivative chromosome consistent with HCT116 haplotype reconstruction by refLinker. Lost segments are indicated by gray-filled boxes.

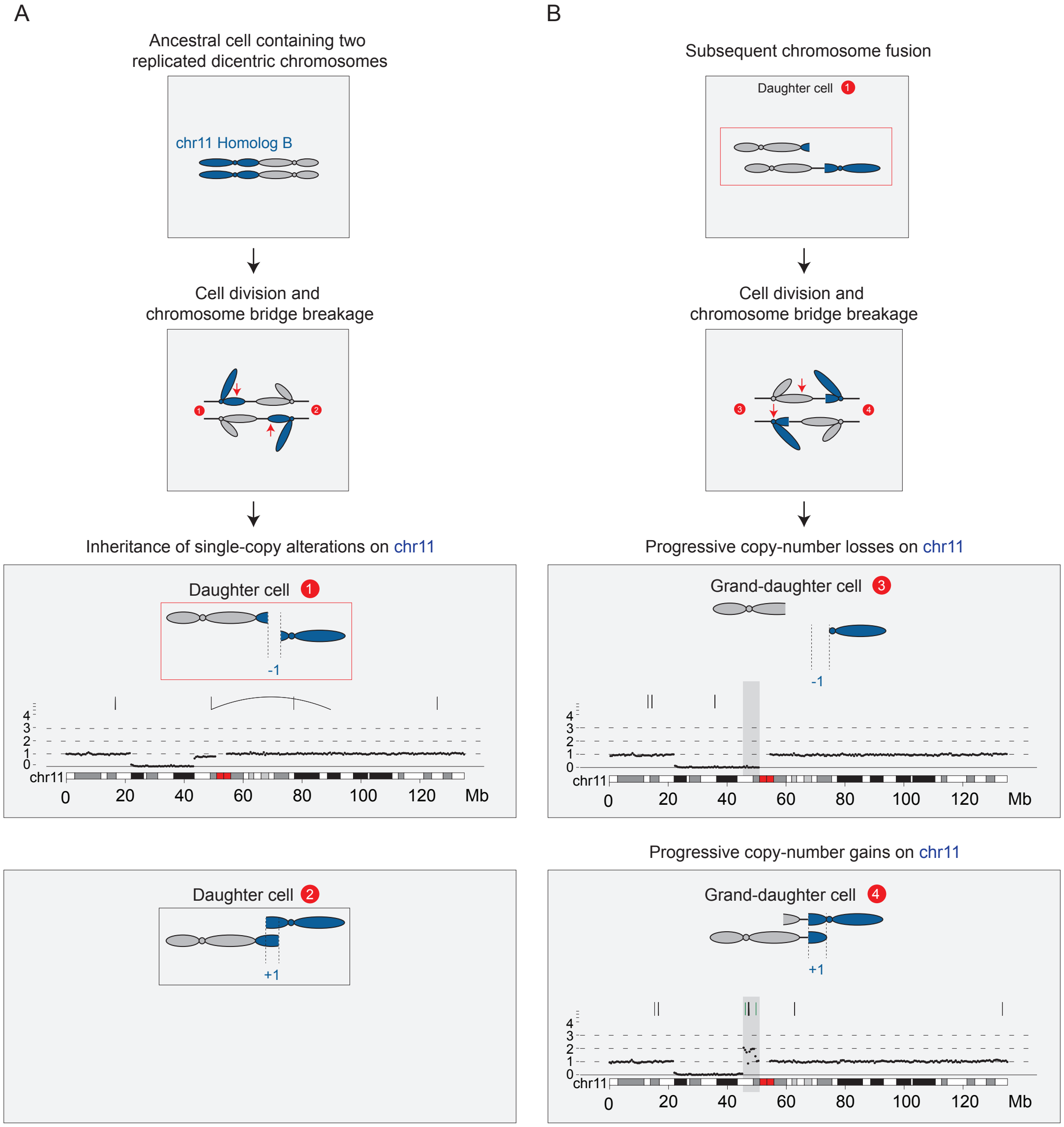


**Fig. S8.** Progressive evolution of single haplotypes following telomere crisis. (A) Top: Schematic showing expected pattern of reciprocal interstitial loss and gain resulting from a single round of replicated dicentric chromosome breakage, where the orientations of segregating dicentric chromosomes are inverted. Bottom: DNA copy number analysis reveals an ancestral interstitial loss on chr11. (B) Top: Subsequent chromosome fusion and breakage in Daughter cell 1 results in progressive reciprocal DNA gains and losses. Bottom: DNA copy-number is plotted for HCT116 sister clones showing reciprocal DNA-copy-number gain and loss adjacent to the ancestral deletion (gray shading). Daughter cell 1: SRR6312277, Grand-daughter cell 3: SRR6312276, Grand-daughter cell 4: SRR6312278.

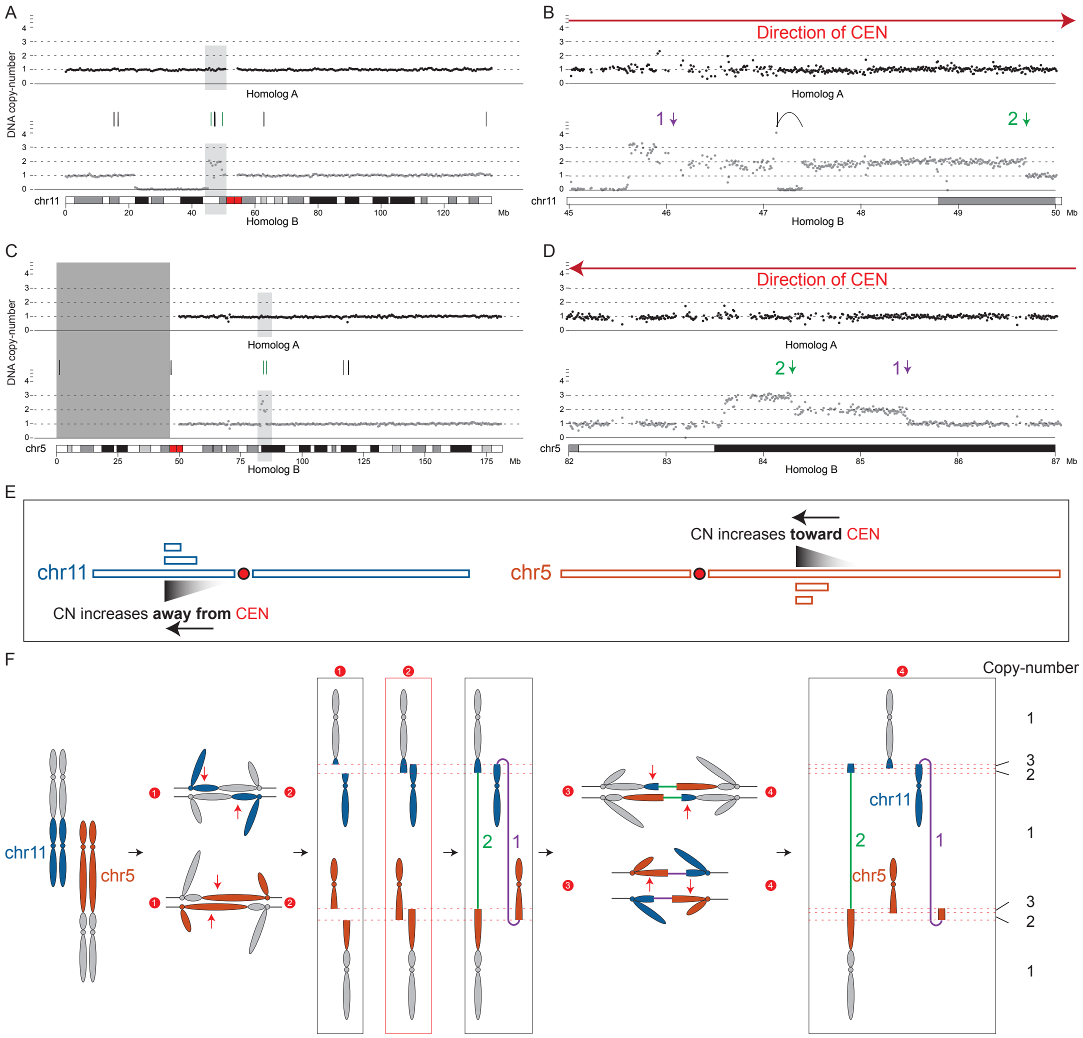


**Fig. S9.** Progressive evolution of single haplotypes following telomere crisis. (A) DNA copy-number is plotted for both chr11 homologs in Grand-daughter cell 4 (500 kb binning). Intrachromosomal translocations are indicated above in black. Interchromosomal breakpoints are indicated in green. Gray shading denotes a five megabase interval spanning a region of intrachromosomal single- and double- DNA copy gains. (B) DNA copy-number is plotted for the 5 Mb gray-shaded region (10 kb binning). Arrows 1 and 2 indicate translocation breakpoints to the corresponding loci in (D). (C) DNA copy-number is plotted as in (A) for both chr5 homologs. Gray shading denotes a five megabase interval spanning a region of intrachromosomal single- and double- DNA copy gains. (D) DNA copy-number is plotted for the 5 Mb gray-shaded region (10 kb binning). Arrows 1 and 2 indicate translocation breakpoints to the corresponding loci in (B). (E) Chromosome schematics showing the orientation of progressive copy-number gains and translocations for chr5B and chr11B. (F) Proposed model for progressive copy-number evolution resulting from the formation of multiple dicentric chromosomes.


Gregory J. Brunette, Richard W. Tourdot, David Pellman, Cheng-Zhong Zhang

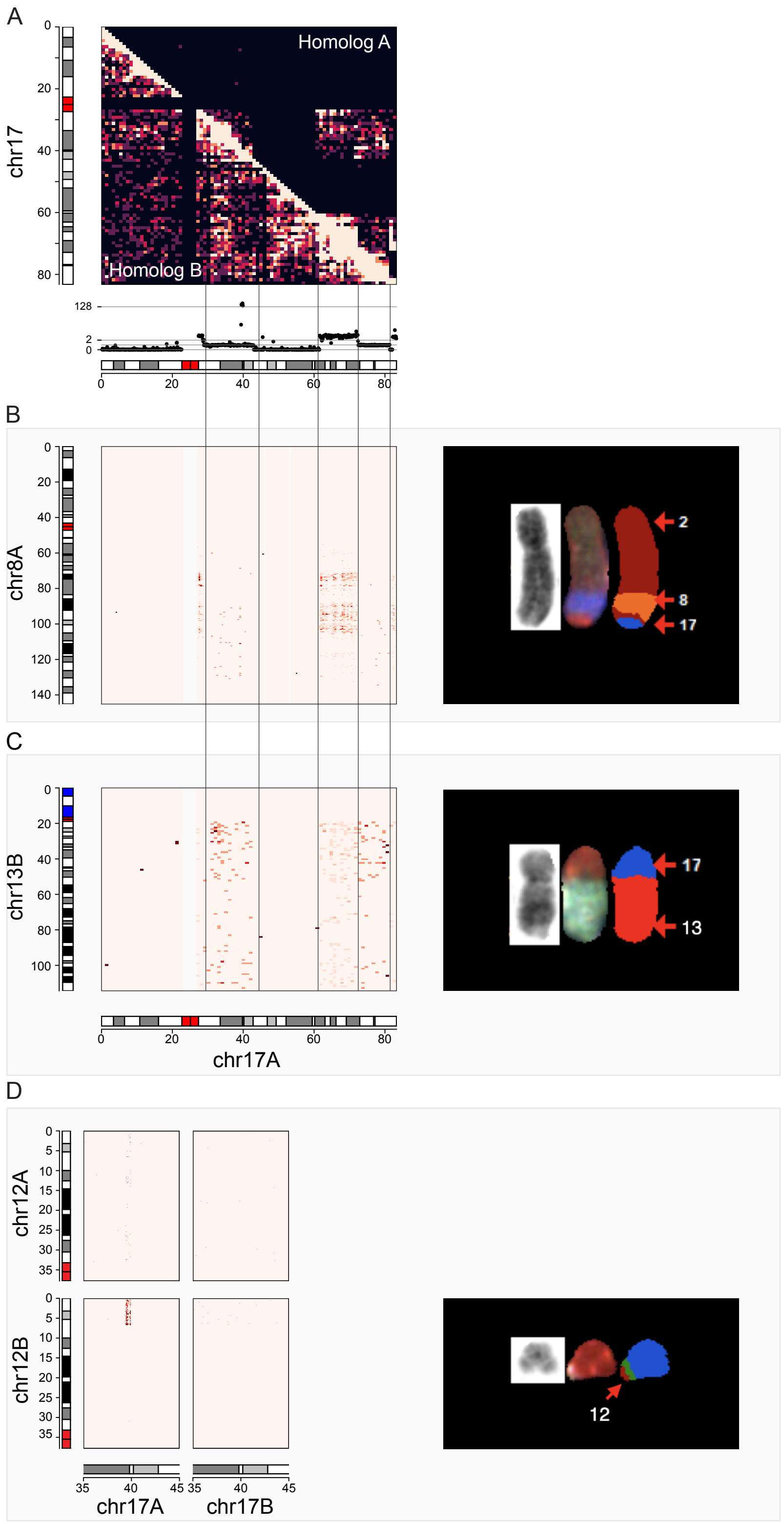


**Fig. S10.** Haplotype-resolved genome complexity in HCC1954. (A) Allele-specific copy-number of HCC1954 chr17. (B) Top: Hi-C maps of HCC1954 chr17 intrachromosomal interactions for homolog A (upper triangle) and homolog B (lower triangle). Bottom: DNA copy number of HCC1954 chr17A, revealing large deletions and high-copy *ERBB2* amplification. (C) Hi-C maps between both chr8 homologs (vertical axis) and chr17 homologs (horizontal axis), revealing chr17A translocation to chr8A. (D) Hi-C maps between both chr13 homologs (vertical axis) and chr17 homologs (horizontal axis) revealing chr17A translocation to chr13B. (E) Hi-C maps placing the *ERBB2* amplicon in a derivative chromosome capped by chr12B telomere. For C, D, and E, Hi-C linkage is normalized by chr17 allelic copy-number.

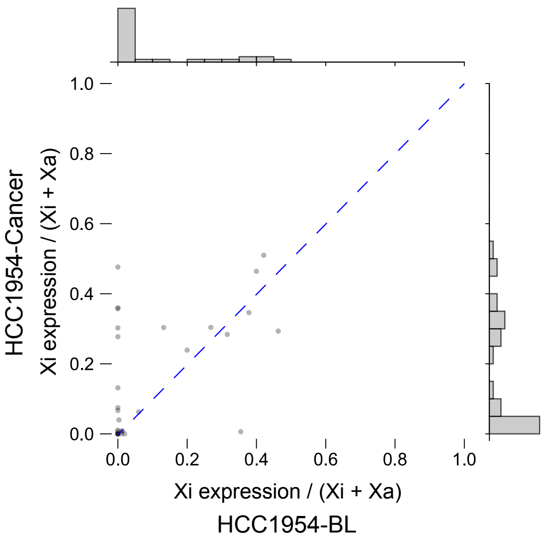


**Fig. S11.** Relative Xi expression for chrXp genes is plotted for HCC1954 (y-axis) versus HCC1954BL (x-axis), a matched normal blood sample with intact X-chromosome inactivation.

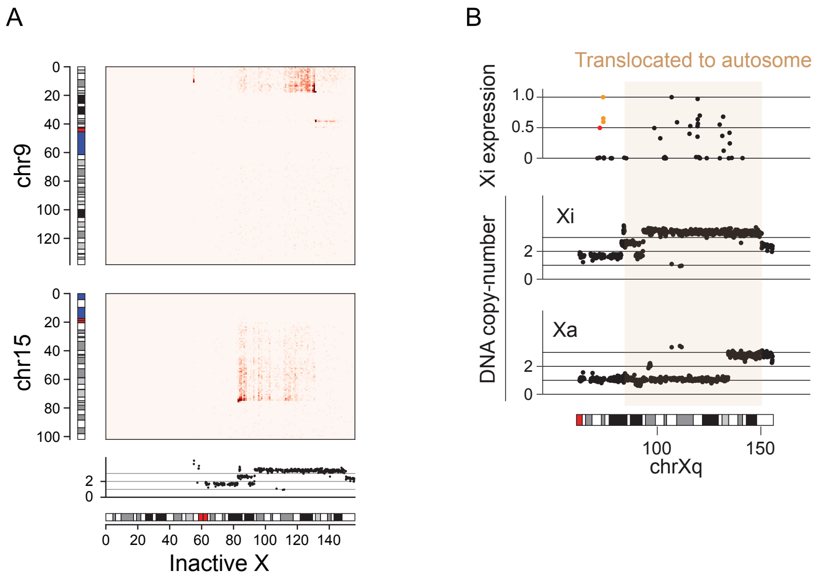


**Fig. S12.** Xi rearrangements and expression in HCC38 (A) Hi-C linkage between chr15B and both chrX alleles supports the chr15-Xi translocation. (B) Allele-specific expression and DNA copy-number of chrXq alleles in HCC38. Top: relative Xi expression is plotted for every gene on HCC38 chrXq. Genes transcribed from the XIC are indicated in orange. A constitutive XCI escapee is plotted in red. Bottom: DNA copy-number is plotted for Xa (lower) and Xi (upper).

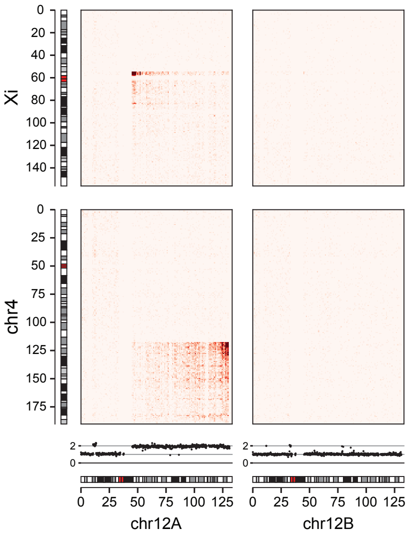


**Fig. S13.** Hi-C reconstruction of Xi-12-4 HCC38 marker chromosome

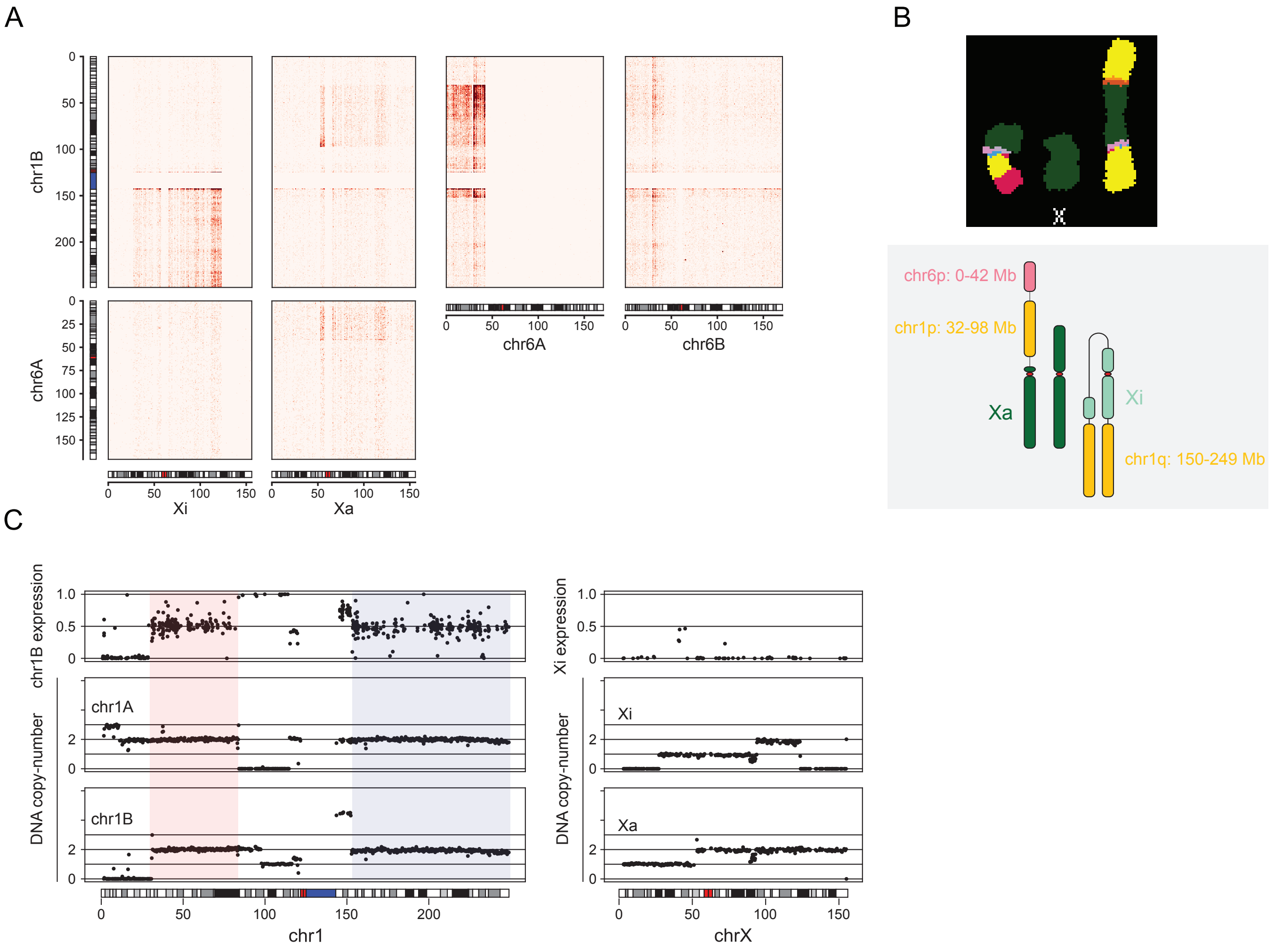


**Fig. S14.** X-linked translocations in HCC1187 (A) Hi-C linkage between chr1B and both chrX alleles supports independent translocations to Xi and Xa. (B) SKY karyotyping (top) and digital schematic (bottom) of tr(1,X) derivative chromosomes in HCC1187. (C) Allele-specific expression and DNA copy-number of chr1 and chrX alleles in HCC1187. Bottom: DNA copy-number is plotted for chr1B (lower) and chr1B (upper). Top: Relative chr1B expression is plotted for every gene on chr1. The p-arm segment translocated to Xa is highlighted in red and the q-arm segment translocated to Xi in blue.

**Table S1. Hi-C data sources**

| Genome | Library size | Long-range contacts[a] | Data source |
|---:|---:|---:|---:|
| NA12878 | 486,848,169[b] | 91,428,507 | (19) |
| RPE-1 | 245,700,771[c] | 29,491,013 | (20) |
| HCC1954BL | 1,255,515,805[d] | 66,804,768 | This study |
| HCC1954 | 1,341,657,004[e] | 91,488,308 | This study |

[a] $> 1$ Mb
[b] SRR1658572: median insert 377; 484,211,662 aligned in pair; 2 ×101bp reads; duplication rate 0.028.
[c] SRR1045724: median insert 364; 193,268,170 aligned in pair; 2 ×150bp reads; duplication rate 0.093.
[d] median insert 308; 1,067,839,232 aligned in pair ; 2 ×150bp reads; duplication rate 0.293.
[e] median insert 248; 842,223,879 aligned in pair; 2 ×250bp reads; duplication rate 0.134.

**Table S2. Genome-wide phasing accuracy**

| Genome | phased in truth set | linked by Hi-C | comparable | accuracy | EAGLE2 recovery | comparable | accuracy |
|---|---|---|---|---|---|---|---|
| NA12878 | 2,156,346 | 1,650,419 | 1,611,090 | 0.997 | 2,043,936 | 1,784,694 | 0.997 |
| RPE-1 | 2,151,642 | 1,819,693 | 1,736,800 | 0.992 | 2,484,304 | 2,061,065 | 0.991 |
| HCC1954BL | 2,027,293 | 2,156,067 | 1,863,261 | 0.993 | 2,459,343 | 1,881,245 | 0.992 |
| HCC1954 | 2,027,293 | 2,010,935 | 1,739,987 | 0.994 | 2,459,230 | 1,881,245 | 0.994 |

**Table S3. refLinker phasing performance for NA12878 data**

| chrom | linked by Hi-C | comparable | accuracy | recovered | comparable | accuracy |
|---|---|---|---|---|---|---|
| chr1 | 131200 | 128946 | 0.997 | 160741 | 141053 | 0.996 |
| chr2 | 131448 | 129036 | 0.997 | 162513 | 142757 | 0.997 |
| chr3 | 120441 | 115992 | 0.997 | 150105 | 129008 | 0.997 |
| chr4 | 100722 | 99225 | 0.997 | 127858 | 112464 | 0.997 |
| chr5 | 104119 | 102225 | 0.997 | 130502 | 114268 | 0.997 |
| chr6 | 117294 | 115371 | 0.997 | 146688 | 128485 | 0.997 |
| chr7 | 96169 | 94489 | 0.997 | 118940 | 104852 | 0.997 |
| chr8 | 84177 | 82553 | 0.998 | 104285 | 91835 | 0.998 |
| chr9 | 74724 | 73640 | 0.997 | 91962 | 81672 | 0.997 |
| chr10 | 86607 | 85348 | 0.994 | 106686 | 94121 | 0.993 |
| chr11 | 83793 | 81048 | 0.998 | 103362 | 89472 | 0.997 |
| chr12 | 75227 | 73738 | 0.998 | 93339 | 81277 | 0.998 |
| chr13 | 62502 | 61826 | 0.997 | 78851 | 69659 | 0.996 |
| chr14 | 56739 | 55806 | 0.998 | 70387 | 61815 | 0.998 |
| chr15 | 47043 | 46176 | 0.997 | 57198 | 50311 | 0.996 |
| chr16 | 33282 | 32564 | 0.997 | 40196 | 35629 | 0.997 |
| chr17 | 41720 | 40557 | 0.995 | 50621 | 43653 | 0.995 |
| chr18 | 38653 | 37924 | 0.996 | 48373 | 42597 | 0.996 |
| chr19 | 35854 | 30121 | 0.990 | 43059 | 31943 | 0.990 |
| chr20 | 38548 | 37849 | 0.997 | 46586 | 41217 | 0.997 |
| chr21 | 23699 | 23431 | 0.997 | 29599 | 26283 | 0.997 |
| chr22 | 22376 | 22007 | 0.996 | 26831 | 23591 | 0.996 |
| chrX | 44082 | 41218 | 0.995 | 55254 | 46732 | 0.994 |
| Total | 1650419 | 1611090 | 0.997 | 2043936 | 1784694 | 0.997 |

**Table S4. refLinker phasing performance for RPE-1 data**

| chrom | linked by Hi-C | comparable | accuracy | recovered | comparable | accuracy |
|---|---|---|---|---|---|---|
| chr1 | 136354 | 131633 | 0.994 | 185675 | 155070 | 0.993 |
| chr2 | 146515 | 141987 | 0.994 | 200686 | 169093 | 0.993 |
| chr3 | 127020 | 117696 | 0.994 | 173387 | 139586 | 0.993 |
| chr4 | 128281 | 124385 | 0.988 | 176931 | 149627 | 0.987 |
| chr5 | 111694 | 107645 | 0.993 | 153652 | 128374 | 0.993 |
| chr6 | 120839 | 115934 | 0.994 | 165256 | 137919 | 0.994 |
| chr7 | 104643 | 101035 | 0.991 | 143611 | 121169 | 0.990 |
| chr8 | 95679 | 92543 | 0.994 | 129656 | 110583 | 0.994 |
| chr9 | 79355 | 76003 | 0.994 | 107999 | 90320 | 0.993 |
| chr10 | 89022 | 85337 | 0.991 | 119615 | 99749 | 0.990 |
| chr11 | 86924 | 82808 | 0.983 | 118252 | 98160 | 0.982 |
| chr12 | 81740 | 78057 | 0.993 | 111077 | 91553 | 0.993 |
| chr13 | 65285 | 63702 | 0.993 | 90596 | 76699 | 0.993 |
| chr14 | 55764 | 53917 | 0.99 | 76702 | 64071 | 0.988 |
| chr15 | 51688 | 50026 | 0.986 | 70392 | 58856 | 0.985 |
| chr16 | 57105 | 54736 | 0.991 | 76727 | 64365 | 0.991 |
| chr17 | 53092 | 50715 | 0.986 | 71675 | 58547 | 0.984 |
| chr18 | 47895 | 46776 | 0.992 | 66108 | 56399 | 0.992 |
| chr19 | 43864 | 34043 | 0.985 | 58872 | 38402 | 0.984 |
| chr20 | 38779 | 37387 | 0.993 | 51906 | 43859 | 0.992 |
| chr21 | 26668 | 25441 | 0.993 | 37151 | 30717 | 0.992 |
| chr22 | 24534 | 22984 | 0.993 | 33245 | 26922 | 0.992 |
| chrX | 46953 | 42010 | 0.990 | 65133 | 51025 | 0.990 |
| Total | 1819693 | 1736800 | 0.992 | 2484304 | 2061065 | 0.991 |

**Table S5. refLinker phasing performance for HCC1954BL data**

| chrom | linked by Hi-C | comparable | accuracy | recovered | comparable | accuracy |
|---|---|---|---|---|---|---|
| chr1 | 163573 | 139086 | 0.995 | 186611 | 140335 | 0.995 |
| chr2 | 175772 | 150266 | 0.994 | 200885 | 151646 | 0.994 |
| chr3 | 149061 | 130897 | 0.996 | 170166 | 131986 | 0.996 |
| chr4 | 152273 | 136205 | 0.997 | 174043 | 137425 | 0.997 |
| chr5 | 130279 | 114596 | 0.995 | 148323 | 115605 | 0.995 |
| chr6 | 141059 | 126065 | 0.99 | 162772 | 128242 | 0.989 |
| chr7 | 126596 | 110396 | 0.99 | 144436 | 111376 | 0.990 |
| chr8 | 115800 | 102100 | 0.993 | 129996 | 103149 | 0.993 |
| chr9 | 85970 | 73382 | 0.984 | 97777 | 74077 | 0.984 |
| chr10 | 112179 | 98831 | 0.994 | 127860 | 99687 | 0.994 |
| chr11 | 108490 | 94058 | 0.99 | 123409 | 94846 | 0.990 |
| chr12 | 97142 | 84047 | 0.995 | 111253 | 84801 | 0.995 |
| chr13 | 74213 | 65743 | 0.997 | 85077 | 66306 | 0.997 |
| chr14 | 68628 | 59533 | 0.975 | 78433 | 60080 | 0.974 |
| chr15 | 59600 | 50428 | 0.994 | 68286 | 50890 | 0.994 |
| chr16 | 69488 | 61133 | 0.994 | 78833 | 61703 | 0.994 |
| chr17 | 55306 | 44157 | 0.989 | 63384 | 44588 | 0.989 |
| chr18 | 60432 | 53223 | 0.992 | 68609 | 53688 | 0.992 |
| chr19 | 41563 | 34504 | 0.990 | 47951 | 34824 | 0.990 |
| chr20 | 46161 | 38584 | 0.991 | 52320 | 38919 | 0.991 |
| chr21 | 29545 | 26383 | 0.995 | 33798 | 26620 | 0.995 |
| chr22 | 30938 | 26428 | 0.992 | 35349 | 26682 | 0.992 |
| chrX | 61999 | 43216 | 0.994 | 69772 | 43770 | 0.994 |
| Total | 2156067 | 1863261 | 0.993 | 2459343 | 1881245 | 0.992 |

**Table S6. refLinker phasing performance for $0.333\times$ down-sampled RPE-1 data**

| chrom | linked by Hi-C | comparable | accuracy | recovered | comparable | accuracy |
|---|---|---|---|---|---|---|
| chr1 | 112897 | 109845 | 0.985 | 185321 | 155067 | 0.982 |
| chr2 | 120897 | 118072 | 0.987 | 200282 | 169092 | 0.984 |
| chr3 | 105647 | 98657 | 0.989 | 173146 | 139586 | 0.986 |
| chr4 | 105628 | 103226 | 0.985 | 176552 | 149626 | 0.982 |
| chr5 | 92111 | 89407 | 0.989 | 153430 | 128373 | 0.986 |
| chr6 | 100356 | 96888 | 0.991 | 165071 | 137917 | 0.989 |
| chr7 | 85906 | 83536 | 0.984 | 143363 | 121165 | 0.981 |
| chr8 | 79293 | 77233 | 0.990 | 129456 | 110583 | 0.988 |
| chr9 | 65203 | 62958 | 0.982 | 107858 | 90314 | 0.978 |
| chr10 | 74588 | 72192 | 0.986 | 119428 | 99739 | 0.983 |
| chr11 | 71770 | 68927 | 0.978 | 118073 | 98160 | 0.975 |
| chr12 | 68948 | 66307 | 0.987 | 110927 | 91553 | 0.985 |
| chr13 | 54235 | 53255 | 0.99 | 90464 | 76698 | 0.989 |
| chr14 | 46135 | 44939 | 0.979 | 76597 | 64070 | 0.977 |
| chr15 | 42677 | 41604 | 0.979 | 70237 | 58855 | 0.977 |
| chr16 | 47027 | 45390 | 0.983 | 76641 | 64365 | 0.980 |
| chr17 | 44236 | 42504 | 0.972 | 71566 | 58547 | 0.968 |
| chr18 | 39425 | 38729 | 0.985 | 66011 | 56398 | 0.982 |
| chr19 | 35965 | 28587 | 0.965 | 58837 | 38402 | 0.961 |
| chr20 | 32070 | 31158 | 0.985 | 51836 | 43859 | 0.982 |
| chr21 | 21891 | 21074 | 0.986 | 37110 | 30717 | 0.984 |
| chr22 | 20107 | 18996 | 0.975 | 33165 | 26922 | 0.973 |
| chrX | 37124 | 33985 | 0.955 | 64762 | 51024 | 0.954 |
| Total | 1504136 | 1447469 | 0.984 | 2480133 | 2061032 | 0.981 |

**Table S7. refLinker phasing performance for $0.167\times$ down-sampled RPE-1 data**

| chrom | linked by Hi-C | comparable | accuracy | recovered | comparable | accuracy |
|---|---|---|---|---|---|---|
| chr1 | 90299 | 87897 | 0.963 | 185193 | 155064 | 0.96 |
| chr2 | 96664 | 94454 | 0.972 | 200209 | 169089 | 0.969 |
| chr3 | 84437 | 78903 | 0.975 | 173046 | 139582 | 0.973 |
| chr4 | 84334 | 82484 | 0.968 | 176433 | 149625 | 0.965 |
| chr5 | 73255 | 71153 | 0.976 | 153378 | 128373 | 0.975 |
| chr6 | 80153 | 77515 | 0.980 | 165000 | 137916 | 0.977 |
| chr7 | 68024 | 66178 | 0.940 | 143318 | 121164 | 0.936 |
| chr8 | 63309 | 61692 | 0.974 | 129405 | 110583 | 0.971 |
| chr9 | 51751 | 50011 | 0.964 | 107750 | 90315 | 0.960 |
| chr10 | 61069 | 59260 | 0.974 | 119363 | 99747 | 0.969 |
| chr11 | 56796 | 54647 | 0.963 | 118008 | 98159 | 0.958 |
| chr12 | 56202 | 54092 | 0.974 | 110824 | 91550 | 0.970 |
| chr13 | 43217 | 42455 | 0.969 | 90452 | 76698 | 0.966 |
| chr14 | 36579 | 35644 | 0.96 | 76577 | 64070 | 0.956 |
| chr15 | 34009 | 33171 | 0.953 | 70212 | 58855 | 0.949 |
| chr16 | 36817 | 35542 | 0.934 | 76615 | 64364 | 0.933 |
| chr17 | 6324 | 5970 | 0.924 | 12623 | 10224 | 0.919 |
| chr18 | 31085 | 30527 | 0.959 | 65982 | 56396 | 0.956 |
| chr19 | 28071 | 22458 | 0.858 | 58818 | 38402 | 0.854 |
| chr20 | 25403 | 24676 | 0.956 | 51816 | 43859 | 0.952 |
| chr21 | 17184 | 16586 | 0.969 | 37072 | 30717 | 0.965 |
| chr22 | 15862 | 15001 | 0.928 | 33160 | 26920 | 0.926 |
| chrX | 28664 | 26196 | 0.625 | 64593 | 51023 | 0.638 |
| Total | 1169508 | 1126512 | 0.955 | 2419847 | 2012695 | 0.952 |

**Table S8. IntegratedPhasing performance for RPE-1 data**

| | Full data | | | $0.333\times$ | | |
|---|---|---|---|---|---|---|
| chrom | phased sites | comparable | accuracy | phased sites | comparable | accuracy |
| chr1 | 148735 | 144936 | 0.981 | 147339 | 143727 | 0.946 |
| chr2 | 163834 | 160325 | 0.990 | 162604 | 159207 | 0.940 |
| chr3 | 136832 | 133770 | 0.989 | 135751 | 132799 | 0.972 |
| chr4 | 144980 | 142267 | 0.985 | 143873 | 141257 | 0.963 |
| chr5 | 123953 | 121230 | 0.988 | 122904 | 120286 | 0.975 |
| chr6 | 133630 | 131206 | 0.992 | 132468 | 130167 | 0.977 |
| chr7 | 115828 | 112936 | 0.988 | 114876 | 112079 | 0.964 |
| chr8 | 107292 | 104946 | 0.991 | 106475 | 104213 | 0.971 |
| chr9 | 87145 | 84607 | 0.988 | 86176 | 83735 | 0.934 |
| chr10 | 95146 | 92984 | 0.988 | 94482 | 92401 | 0.961 |
| chr11 | 93603 | 91339 | 0.982 | 93033 | 90843 | 0.958 |
| chr12 | 90148 | 87407 | 0.992 | 89629 | 86995 | 0.947 |
| chr13 | 74410 | 73314 | 0.991 | 73810 | 72766 | 0.976 |
| chr14 | 60981 | 59514 | 0.990 | 60502 | 59078 | 0.951 |
| chr15 | 56569 | 55371 | 0.978 | 56090 | 54943 | 0.967 |
| chr16 | 63475 | 61582 | 0.989 | 62966 | 61152 | 0.963 |
| chr17 | 54584 | 52643 | 0.972 | 54096 | 52224 | 0.948 |
| chr18 | 55223 | 54244 | 0.990 | 54946 | 54010 | 0.975 |
| chr19 | 37601 | 36011 | 0.975 | 37320 | 35808 | 0.926 |
| chr20 | 42454 | 41462 | 0.988 | 42057 | 41114 | 0.956 |
| chr21 | 29865 | 29388 | 0.993 | 29674 | 29216 | 0.981 |
| chr22 | 26119 | 25343 | 0.981 | 25970 | 25216 | 0.924 |
| Total | 1942407 | 1896825 | 0.987 | 1927041 | 1883236 | 0.959 |

**Table S9. refLinker phasing performance for HCC1954 (cancer) data**

| chrom | linked by Hi-C | comparable | accuracy | recovered | comparable | accuracy |
|---|---|---|---|---|---|---|
| chr1 | 162285 | 137968 | 0.997 | 186611 | 140335 | 0.997 |
| chr2 | 174356 | 149071 | 0.996 | 200884 | 151646 | 0.996 |
| chr3 | 147570 | 129571 | 0.998 | 170167 | 131986 | 0.997 |
| chr4 | 150847 | 134871 | 0.998 | 174043 | 137425 | 0.998 |
| chr5 | 79511 | 70078 | 0.995 | 148296 | 115605 | 0.995 |
| chr6 | 140938 | 126004 | 0.991 | 162773 | 128242 | 0.991 |
| chr7 | 125878 | 109738 | 0.993 | 144436 | 111376 | 0.993 |
| chr8 | 86317 | 75662 | 0.992 | 129988 | 103149 | 0.990 |
| chr9 | 84708 | 72322 | 0.987 | 97773 | 74077 | 0.986 |
| chr10 | 111005 | 97792 | 0.995 | 127860 | 99687 | 0.995 |
| chr11 | 107447 | 93099 | 0.992 | 123409 | 94846 | 0.992 |
| chr12 | 86795 | 74693 | 0.996 | 111250 | 84801 | 0.995 |
| chr13 | 73596 | 65187 | 0.999 | 85077 | 66306 | 0.998 |
| chr14 | 67715 | 58688 | 0.976 | 78432 | 60080 | 0.976 |
| chr15 | 59227 | 50117 | 0.997 | 68285 | 50890 | 0.996 |
| chr16 | 69173 | 60890 | 0.996 | 78833 | 61703 | 0.996 |
| chr17 | 41902 | 33600 | 0.99 | 63374 | 44588 | 0.991 |
| chr18 | 59526 | 52429 | 0.993 | 68608 | 53688 | 0.993 |
| chr19 | 39280 | 32564 | 0.994 | 47951 | 34824 | 0.993 |
| chr20 | 45835 | 38309 | 0.995 | 52320 | 38919 | 0.994 |
| chr21 | 29028 | 25901 | 0.998 | 33796 | 26620 | 0.998 |
| chr22 | 26906 | 23021 | 0.995 | 35346 | 26682 | 0.994 |
| chrX | 41090 | 28412 | 0.995 | 69718 | 43770 | 0.994 |
| Total | 2010935 | 1739987 | 0.994 | 2459230 | 1881245 | 0.994 |

**SI Dataset S1 (HCT116_refLinker_haplotype_chr{1..22}.dat)**

refLinker-generated haplotypes for the HCT116 cell line can be found at https://github.com/gbrunette/refLinker/tree/main/haplotype_data.

**SI Dataset S2 (HCC1954_refLinker_haplotype_chr{1..22}X.dat)**

refLinker-generated haplotypes for the HCC1954 cell line can be found at https://github.com/gbrunette/refLinker/tree/main/haplotype_data
